# Sperm membrane proteins DCST1 and DCST2 are required for the sperm-egg fusion process in mice and fish

**DOI:** 10.1101/2021.04.18.440256

**Authors:** Taichi Noda, Andreas Blaha, Yoshitaka Fujihara, Krista R. Gert, Chihiro Emori, Victoria E. Deneke, Seiya Oura, Sara Berent, Mayo Kodani, Karin Panser, Luis Enrique Cabrera-Quio, Andrea Pauli, Masahito Ikawa

**Affiliations:** Research Institute for Microbial Diseases, Osaka University, 3-1 Yamadaoka, Suita, Osaka 565-0871, Japan; Institute of Resource Development and Analysis, Kumamoto University, 2-2-1 Honjo, Kumamoto 860-0811, Japan; Research Institute of Molecular Pathology (IMP), Vienna BioCenter (VBC), Campus-Vienna-Biocenter 1, 1030 Vienna, Austria; Vienna BioCenter PhD Program, Doctoral School of the University at Vienna and Medical University of Vienna, Vienna, Austria; Department of Bioscience and Genetics, National Cerebral and Cardiovascular Center, 6-1 Kishibe-Shimmachi, Suita, Osaka 564-8565, Japan; Graduate School of Pharmaceutical Sciences, Osaka University, 1-6 Yamadaoka, Suita, Osaka 565-0871, Japan; The Institute of Medical Science, The University of Tokyo, 4-6-1 Shirokanedai, Minato-ku, Tokyo 108-8639, Japan

## Abstract

The process of sperm-egg fusion is critical for successful fertilization, yet the underlying mechanisms that regulate these steps have remained unclear in vertebrates. Here, we show that both mouse and zebrafish DCST1 and DCST2 are necessary in sperm to fertilize the egg, similar to their orthologs SPE-42 and SPE-49 in *C. elegans* and Sneaky in *D. melanogaster*. Mouse *Dcst1* and *Dcst2* single knockout (KO) spermatozoa are able to undergo the acrosome reaction and show normal relocalization of IZUMO1, an essential factor for sperm-egg fusion, to the equatorial segment. While both single KO spermatozoa can bind to the oolemma, they rarely fuse with oocytes, resulting in male sterility. Similar to mice, zebrafish *dcst1* KO males are subfertile and *dcst2* and *dcst1/2* double KO males are sterile. Zebrafish *dcst1/2* KO spermatozoa are motile and can approach the egg, but rarely bind to the oolemma. These data demonstrate that DCST1/2 are essential for male fertility in two vertebrate species, highlighting their crucial role as conserved factors in fertilization.

## Introduction

Until recently, only a few factors had been shown to be essential for the sperm-egg fusion process: IZUMO1 on the sperm membrane and its receptor (IZUMO1R, also known as JUNO and FOLR4) on the egg membrane (oolemma)^1, 2^. Mammalian IZUMO1 and JUNO form a 1:1 complex which is necessary for sperm-egg adhesion prior to fusion^1, 3, 4^. Furthermore, egg-expressed CD9 is also required for sperm-egg fusion, yet its role appears to be indirect by regulating microvilli formation on the oolemma rather than fusion^5–8^. Recently, we and other research groups have found that four additional sperm factors [fertilization influencing membrane protein (FIMP), sperm-oocyte fusion required 1 (SOF1), transmembrane protein 95 (TMEM95), and sperm acrosome associated 6 (SPACA6)] are also essential for the sperm-egg fusion process and male fertility in mice^9–12^. However, HEK293 cells expressing all of these sperm-expressed, fusion-related factors in addition to IZUMO1 were able to bind but not fuse with zona pellucida (ZP)-free eggs, suggesting that additional fusion-related factors are necessary for the completion of sperm-egg fusion^9^.

DCSTAMP and OCSTAMP proteins represent an interesting group of proteins to study in the context of cell-cell fusion, since they have been shown to play a role in osteoclast and foreign body giant cell (FBGC) fusion^13–15^. They belong to the class of DC-STAMP-like domain-containing proteins and are multi-pass transmembrane proteins with an intracellular C-terminus containing a non-canonical RING finger domain^13, 16, 17^. DCSTAMP was shown to localize to the plasma membrane and endoplasmic reticulum (ER) membrane in dendritic cells and osteoclasts^16–19^. These cell types in *Dcstamp* KO mice show no apparent defect in differentiation into the osteoclast lineage and cytoskeletal structure, yet osteoclasts and FBGCs are unable to fuse to form terminally differentiated multinucleated cells^14^. Even though OCSTAMP is widely expressed in mouse tissues^20^, the only reported defect in *Ocstamp* KO mice is the inability to form multinucleated osteoclasts and FBGCs^13, 15^. The fusion defect is not due to a change in the expression levels of osteoclast markers, including *Dcstamp*^13, 15^. These results established an essential role for DC-STAMP-like domain-containing proteins in cell-cell fusion.

DC-STAMP-like domain-containing proteins, namely testis-enriched Sneaky, SPE-42, and SPE-49, are necessary for male fertility in *Drosophila*^21, 22^ and *C. elegans*^23–25^, respectively. Specifically, *sneaky*-disrupted fly spermatozoa can enter the egg, but fail to break down the sperm plasma membrane; the male pronucleus thus does not form and embryonic mitotic divisions do not occur^22^. *Spe-42* and *spe-49* mutant *C. elegans* spermatozoa can migrate into the spermatheca, the site of fertilization in worms, but these mutants are nearly or completely sterile, respectively, suggesting that SPE-42 and SPE-49 are involved in the ability of spermatozoa to fertilize eggs^23–25^. Sneaky, SPE-42 and SPE-49 have homologs in vertebrates called DCST1 and DCST2, but the roles of these proteins have remained undetermined. Here, we analyzed the physiological function of *Dcst1* and *Dcst2* and their effect on sperm fertility using genetically modified mice and zebrafish.

## Results

### DCST1 and DCST2 belong to a conserved group of DC-STAMP-like domain-containing proteins

DC-STAMP-like domain-containing proteins are conserved in metazoa. Phylogenetic analysis revealed a split between the orthologous groups of DCSTAMP and OCSTAMP and of DCST1 and DCST2 (Figure S1A). Furthermore, DCSTAMP and OCSTAMP as well as DCST1 and DCST2 orthologs form distinct clades, suggesting two gene duplication events at their origin.

Consistent with this, protein sequence identity between the mouse and zebrafish orthologs of DCST1 (39.6% identity) and DCST2 (38.3% identity) is higher than the sequence identity between paralogs (mouse DCST1 and mouse DCST2: 22.5%; zebrafish Dcst1 and zebrafish Dcst2: 21.3%) (Figure S1B). Based on transmembrane predictions using TMHMM and Phobius^26, 27^, mouse and zebrafish DCST1 and DCST2 (DCST1/2) have five or six transmembrane helices (Figure S1C). Their intracellular C-termini contain six invariant cysteines that are thought to form a non-canonical RING finger domain and are required for SPE-42 function in *C. elegans*^25^. However, the physiological requirements of DCST1/2 in vertebrates have remained unclear.

### DCST1 and DCST2 are required for male fertility in mice

RT-PCR analysis with multiple mouse tissues showed that *Dcst1* and *Dcst2* mRNAs are abundantly expressed in mouse testis (Figure 1A). Using published single-cell RNA-sequencing data^28^, we found that *Dcst1* and *Dcst2* mRNAs peak in mid-round spermatids, indicating that the expression patterns of *Dcst1* and *Dcst2* are similar to that of the other sperm-egg fusion-related genes *Izumo1*, *Fimp*, *Sof1*, *Tmem95*, and *Spaca6* (Figure 1B).

**Figure 1.**
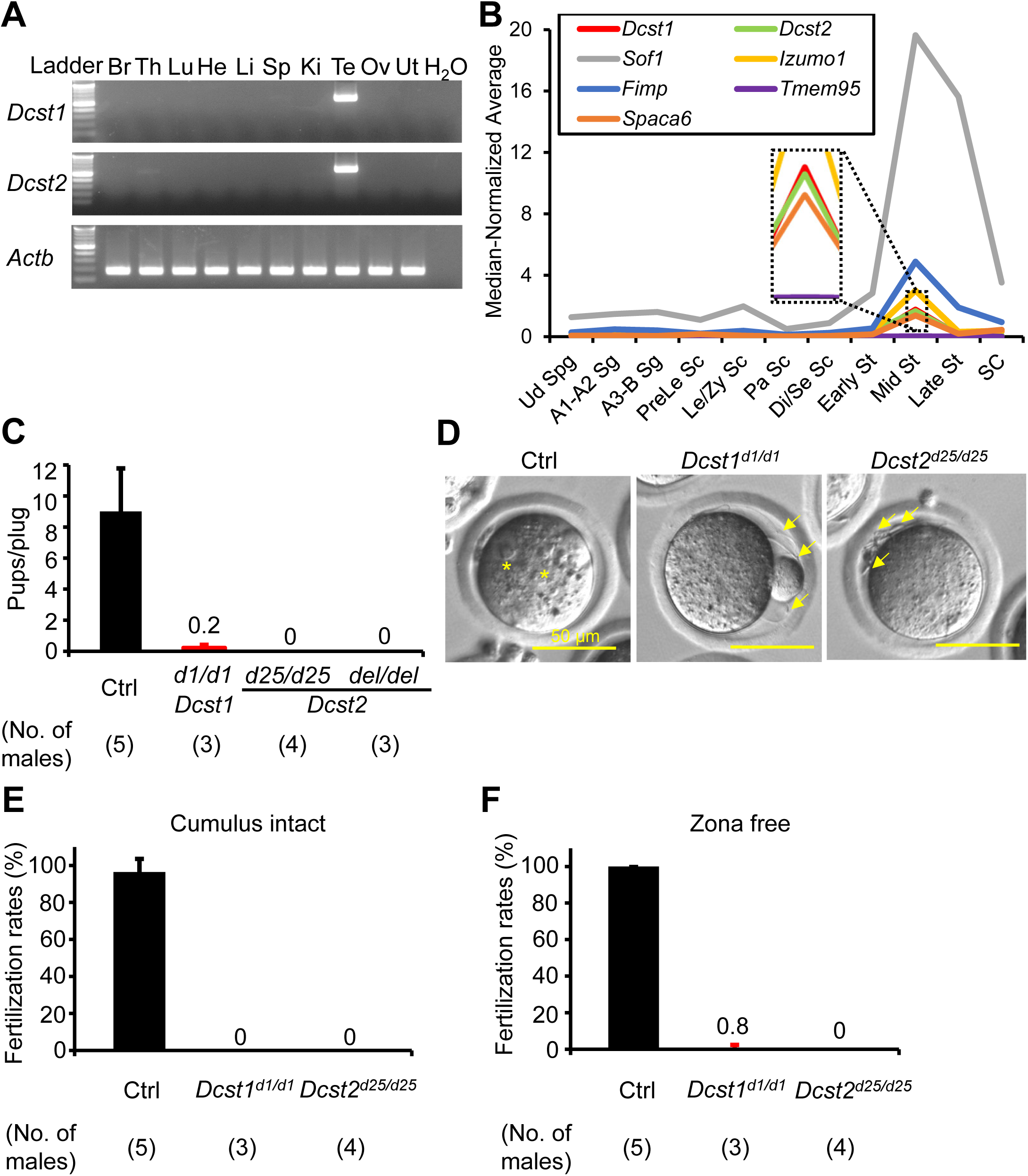
Male fertility of *Dcst1* and *Dcst2* mutant mice. **A) Multi-tissue gene expression analysis.** *Dcst1* and *Dcst2* are abundantly expressed in the mouse testis. Beta actin (*Actb*) was used as the loading control. Br, brain; Th, thymus; Lu, lung; He, heart; Li, liver; Sp, spleen; Ki, kidney; Te, testis; Ov, ovary; Ut, uterus. **B) Median-normalized level of *Dcst1* and *Dcst2* mRNA expression during mouse spermatogenesis.** *Dcst1* and *Dcst2* are strongly expressed in mid-round spermatids, corresponding to other fusion-related factors. Ud Sg, undifferentiated spermatogonia; A1-A2 Sg, A1-A2 differentiating spermatogonia; A3-B Sg, A3-A4-In-B differentiating spermatogonia; Prele Sc, preleptotene spermatocytes; Le/Zy Sc, leptotene/zygotene spermatocytes; Pa Sc, pachytene spermatocytes; Di/Se Sc, diplotene/secondary spermatocytes; Early St, early round spermatids; Mid St, mid round spermatids; Late St, late round spermatids; SC, Sertoli cells. **C) Male fecundity.** Each male was caged with 2 wild-type females for more than 1 month. *Dcst2^d25/wt^* ^and^ *^del^*^/^*^wt^* males were used as the control (Ctrl). *Dcst1^d1/d1^*, *Dcst2^d25/d25^*, and *Dcst2^del/del^* males succeeded in mating [number of plugs: 19 (Ctrl), 17 (*Dcst1^d1/d1^*), 42 (*Dcst2^d25/d25^*), 24 (*Dcst2^del/del^*)], but the females very rarely delivered pups [pups/plug: 9.0 ± 2.8 (Ctrl), 0.2 ± 0.2 (*Dcst1^d1/d1^*), 0 (*Dcst2^d25/d25^*), 0 (*Dcst2^del/del^*)]. **D) Egg observation after IVF.** After 8 hours of incubation, pronuclei were observed in the control spermatozoa (asterisks). However, *Dcst1* KO and *Dcst2* KO spermatozoa accumulated in the perivitelline space (arrows). **E) Sperm fertilizing ability using cumulus-intact eggs *in vitro*.** *Dcst1* KO and *Dcst2* KO spermatozoa could not fertilize eggs [fertilization rates: 96.5 ± 7.1% (Ctrl, 231 eggs), 0% (*Dcst1^d1/d1^*, 97 eggs), 0% (*Dcst2^d25/d25^*, 197 eggs)]. **F) Sperm fertilizing ability using ZP-free eggs *in vitro*.** *Dcst1* KO and *Dcst2* KO spermatozoa rarely fertilized eggs [fertilization rates: 100% (Ctrl, 142 eggs), 0.8 ± 1.6% (*Dcst1^d1/d1^*, 94 eggs), 0% (*Dcst2^d25/d25^*, 88 eggs)].

Using CRISPR/Cas9-mediated mutagenesis, we generated *Dcst2* mutant mice lacking 7,223 bp (*Dcst2^del/del^*), which resulted in the deletion of almost all of the *Dcst2* open reading frame (ORF) (Figure S2A-C). Of note, the expression level of *Dcst1* mRNA in *Dcst2^del/del^* testis decreased (Figure S2D), suggesting that the deleted region is required for *Dcst1* expression in the testis. As shown in Figure S2A, *Dcst1* and *Dcst2* are tandemly arranged such that parts of their 5’ genomic regions overlap. To assess the role of each gene, we generated *Dcst1* indel mice (*Dcst1^d1/d1^*) and *Dcst2* indel mice (*Dcst2^d^*^25^*^/d25^*) (Figure S3A **and** B). RNA isolation from mutant testes followed by cDNA sequencing revealed that *Dcst1^d1/d1^* has a 1-bp deletion in exon 1, and *Dcst2^d25/d25^* has a 25-bp deletion in exon 4 (Figure S3C **and** D). Both deletions result in frameshift mutations leading to premature stop codons.

*Dcst1^d1/d1^*, *Dcst2^d25/d25^*, and *Dcst2^del/del^* male mice successfully mated with female mice, but the females rarely delivered offspring {pups/plug: 9.01 ± 2.77 [control (Ctrl), 19 plugs], 0.22 ± 0.19 [*Dcst1^d1/d1^*, 17 plugs], 0 [*Dcst2^d25/d25^*, 42 plugs], 0 [*Dcst2^del/del^*, 24 plugs]}, indicating that *Dcst1* mutant males are almost and *Dcst2* males are completely sterile (Figure 1C). Unexpectedly, the indel mutations in *Dcst1^d1/d1^* and *Dcst2 ^d25/d25^* decreased the expression level of *Dcst1* mRNA in *Dcst2^d25/d25^* testis and *Dcst2* mRNA in *Dcst1^d1/d1^* testis, respectively (Figure S3C). To evaluate the influence of the decreased expression level of *Dcst1* and *Dcst2* mRNAs on male fertility, we obtained double heterozygous (*Dcst1^d1/wt^* and *Dcst2^del/wt^*) (dHZ) males through intercrossing. DHZ males showed a decreased expression level of both *Dcst1* and *Dcst2* mRNA in the testis, but their fertility was comparable to that of the control (Figure S3E), indicating that the expression levels of *Dcst1* mRNA from the *Dcst2^d25^* allele and *Dcst2* mRNA from the *Dcst1^d1^* allele are decreased but still sufficient to maintain male fertility. This data reconfirms that DCST2 is indispensable for male fertility. Hereafter, we used *Dcst1^d1/d1^* and *Dcst2^d25/d25^* male mice for all experiments unless otherwise specified.

### Spermatozoa from *Dcst1^d1/d1^* and *Dcst2^d25/d25^* mice rarely fertilize eggs

The gross morphology of *Dcst1^d1/d1^* and *Dcst2^d25/d25^* testes was comparable to the control (Figure S4A). Although the testis weight of *Dcst1^d1/d1^* was slightly reduced [testis weight (mg)/body weight (g): 3.13 ± 0.19 *(Dcst1^d1/wt^*), 2.56 ± 0.27 (*Dcst1^d1/d1^*), 3.88 ± 0.34 (*Dcst2^d25/wt^*), 3.60 ± 0.28 (*Dcst2^d25/d25^*)] (Figure S4B), PAS-hematoxylin staining revealed no overt defects in spermatogenesis of *Dcst1^d1/d1^* and *Dcst2^d25/d25^* males (Figure S4C). The sperm morphology and motility parameters of *Dcst1^d1/d1^* and *Dcst2^d25/d25^* mice were normal (Figure S5). However, when the mutant spermatozoa were incubated with cumulus-intact wild-type eggs *in vitro*, they accumulated in the perivitelline space and could not fertilize eggs [96.5 ± 7.1% (Ctrl, 231 eggs), 0% (*Dcst1^d1/d1^*, 97 eggs), and 0% (*Dcst2^d25/d25^*, 197 eggs)] (Figure 1D-E **and Movies S1-2**). Furthermore, even when these KO spermatozoa were incubated with ZP-free eggs, only one egg and no eggs were fertilized by *Dcst1* KO spermatozoa and *Dcst2* KO spermatozoa, respectively [100% (Ctrl, 142 eggs), 0.8 ± 1.6% (*Dcst1^d1/d1^*, 94 eggs), 0% (*Dcst2^d25/d25^*, 88 eggs)] (Figure 1F).

### Spermatozoa from *Dcst1^d1/d1^* and *Dcst2^d25/d25^* mice can bind to, but not fuse with eggs

To examine the binding and fusion ability of *Dcst1^d1/d1^* and *Dcst2^d25/d25^* mutant spermatozoa, we incubated these mutant spermatozoa with ZP-free eggs. Both mutant spermatozoa could bind to the oolemma [5.72 ± 1.97 (Ctrl, 113 eggs), 7.64 ± 4.68 (*Dcst1^d1/d1^*, 89 eggs), 7.63 ± 3.45 (*Dcst2^d25/d25^*, 89 eggs)] (Figure 2A and B). Because binding is not defective in mutant spermatozoa, we confirmed that IZUMO1, a key factor in this process, was expressed and localized normally in testicular germ cells (TGCs) and spermatozoa of *Dcst1^d1/d1^* and *Dcst2^d25/d25^* males (Figure 2C). Indeed, we found that the level of IZUMO1 in mutant spermatozoa was comparable to the control (Figure 2C). Moreover, there was no difference in the acrosome reaction rate of oolemma-bound spermatozoa between control, *Dcst1^d1/d1^* and *Dcst2^d25/d25^* males, determined by live-cell staining with IZUMO1 antibody [58.9 ± 17.9% (Ctrl), 80.5 ± 4.6% (*Dcst1^d1/d1^*), 63.5± 6.5% (*Dcst2^d25/d25^*)] (Figure 2D and E). Next, the mutant spermatozoa were incubated with Hoechst 33342-preloaded ZP-free eggs. In experiments with control spermatozoa, Hoechst 33342 fluorescence signal was translocated into sperm heads (Figure 2F), indicating that these spermatozoa fused with the egg membrane. However, Hoechst 33342 signal was rarely detected in *Dcst1* KO and *Dcst2* KO spermatozoa bound to the oolemma [fused spermatozoa/egg: 1.52 ± 0.35 (Ctrl, 113 eggs), 0.04 ± 0.05 (*Dcst1^d1/d1^*, 73 eggs), 0 (*Dcst2^d25/d25^*, 73 eggs)] (Figure 2F and G). Thus, *Dcst1* and *Dcst2* KO spermatozoa can bind to eggs but not fuse with them.

**Figure 2.**
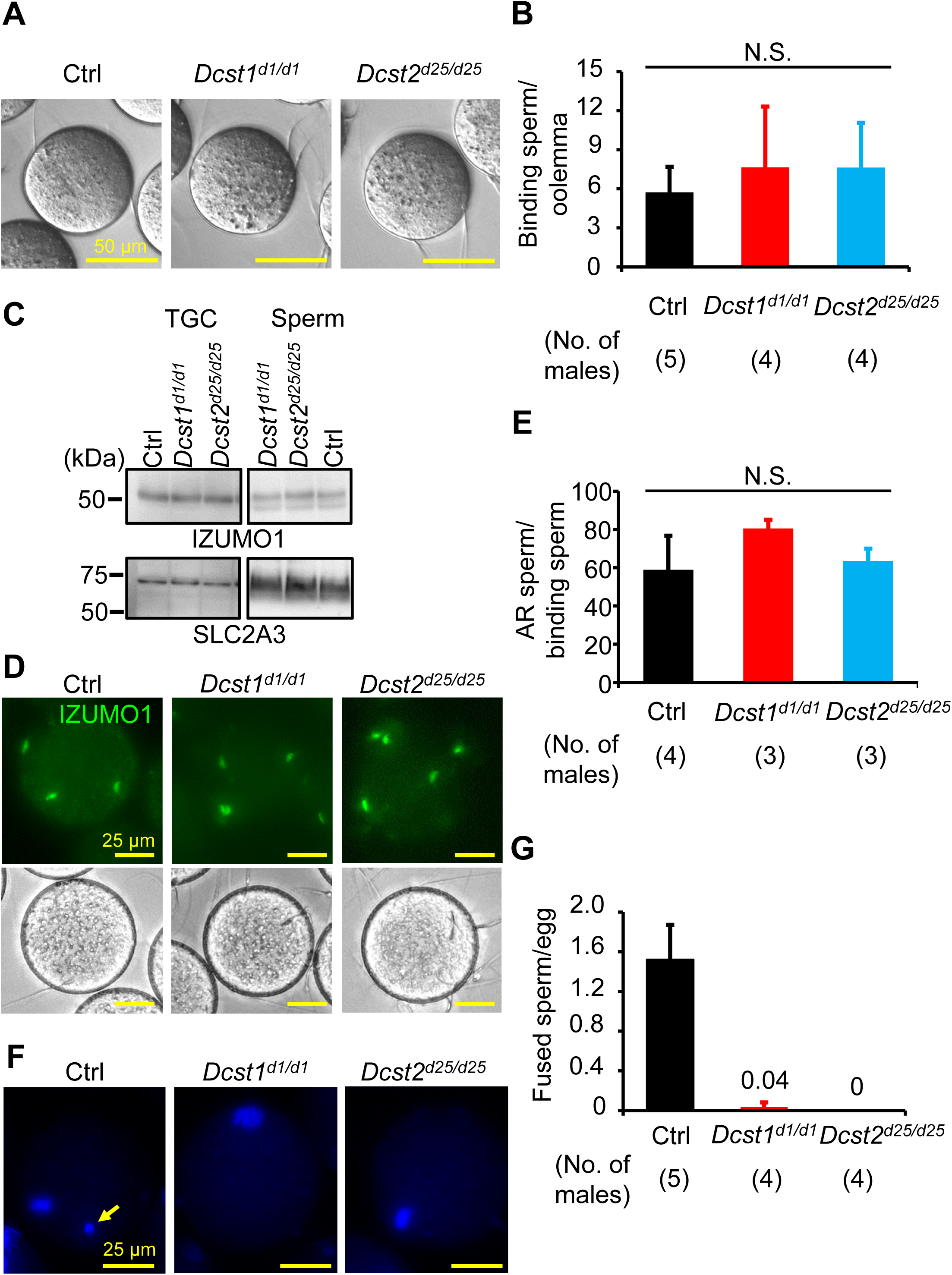
Adhesion and fusion ability of *Dcst1* and *Dcst2* mutant spermatozoa to oocyte plasma membrane. **A and B) Binding ability.** *Dcst1* KO and *Dcst2* KO spermatozoa could bind to the oolemma after 30 minutes of incubation (panel A). There is no significant difference in the sperm number bound to the oolemma (panel B) [binding spermatozoa/egg: 5.7 ± 2.0 (Ctrl, 113 eggs), 7.6 ± 4.7 (*Dcst1^d1/d1^*, 89 eggs), 7.6 ± 3.5 (*Dcst2^d25/d25^*, 89 eggs)]. N.S.: not significant (p > 0.05). **C) Detection of IZUMO1.** The band signals of IZUMO1 in TGC and spermatozoa of *Dcst1^d1/d1^* and *Dcst2^d25/d25^* male mice were comparable to the control. SLC2A3, one of proteins in sperm tail, was used as the loading control. **D and E) Acrosome status of binding spermatozoa.** Live spermatozoa bound to the oolemma were stained with the IZUMO1 antibody, and IZUMO1 only in the acrosome reacted (AR) spermatozoa was detected (panel D). There are no significant differences in the rates of AR spermatozoa (panel E) [AR spermatozoa/binding spermatozoa: 58.9 ± 17.9% (Ctrl), 80.5 ± 4.6% (*Dcst1^d1/d1^*), 63.5 ± 6.5% (*Dcst2^d25/d25^*)]. N.S.: not significant (p = 0.13). **F and G) Fusion ability.** The ZP-free eggs pre-stained Hoechst 33342 were used for sperm-egg fusion assay. Hoechst 33342 signal transferred to control sperm heads, indicating that spermatozoa fused with eggs (panel F, arrow). However, *Dcst1* KO and *Dcst2* KO spermatozoa barely fused with eggs [fused spermatozoa/egg: 1.52 ± 0.35 (Ctrl, 113 eggs), 0.04 ± 0.05 (*Dcst1^d1/d1^*, 73 eggs), 0 (*Dcst2^d25/d25^*, 73 eggs)].

### Sterility of *Dcst1^d1/d1^* and *Dcst2^d25/d25^* males is rescued by *Dcst1*-3xHA and *Dcst2*-3xHA transgenes

To confirm that the *Dcst1* and *Dcst2* disruptions are responsible for male sterility, we generated transgenic mice in which a testis-specific Calmegin (*Clgn*) promoter expresses mouse DCST1 and DCST2 with an HA tag at the C-terminus (Figure S6A **and** B). When *Dcst1^d1/d1^* males with the *Dcst1*-3xHA transgene and *Dcst2^d25/d25^* males with the *Dcst2*-3xHA transgene were mated with wild-type females, the females delivered normal numbers of offspring [pups/plug: 5.7 ± 0.5 (*Dcst1^d1/d1^*; Tg, 25 plugs), 7.6 ± 2.7 (*Dcst2^d25/d25^*; Tg, 15 plugs)] (Figure 3A). We could detect HA-tagged DCST1 and HA-tagged DCST2 in TGCs and spermatozoa at the expected sizes for the full-length proteins (Figure 3B, arrowheads), though both proteins appear to be subject to post-translational processing or protein degradation.

**Figure 3.**
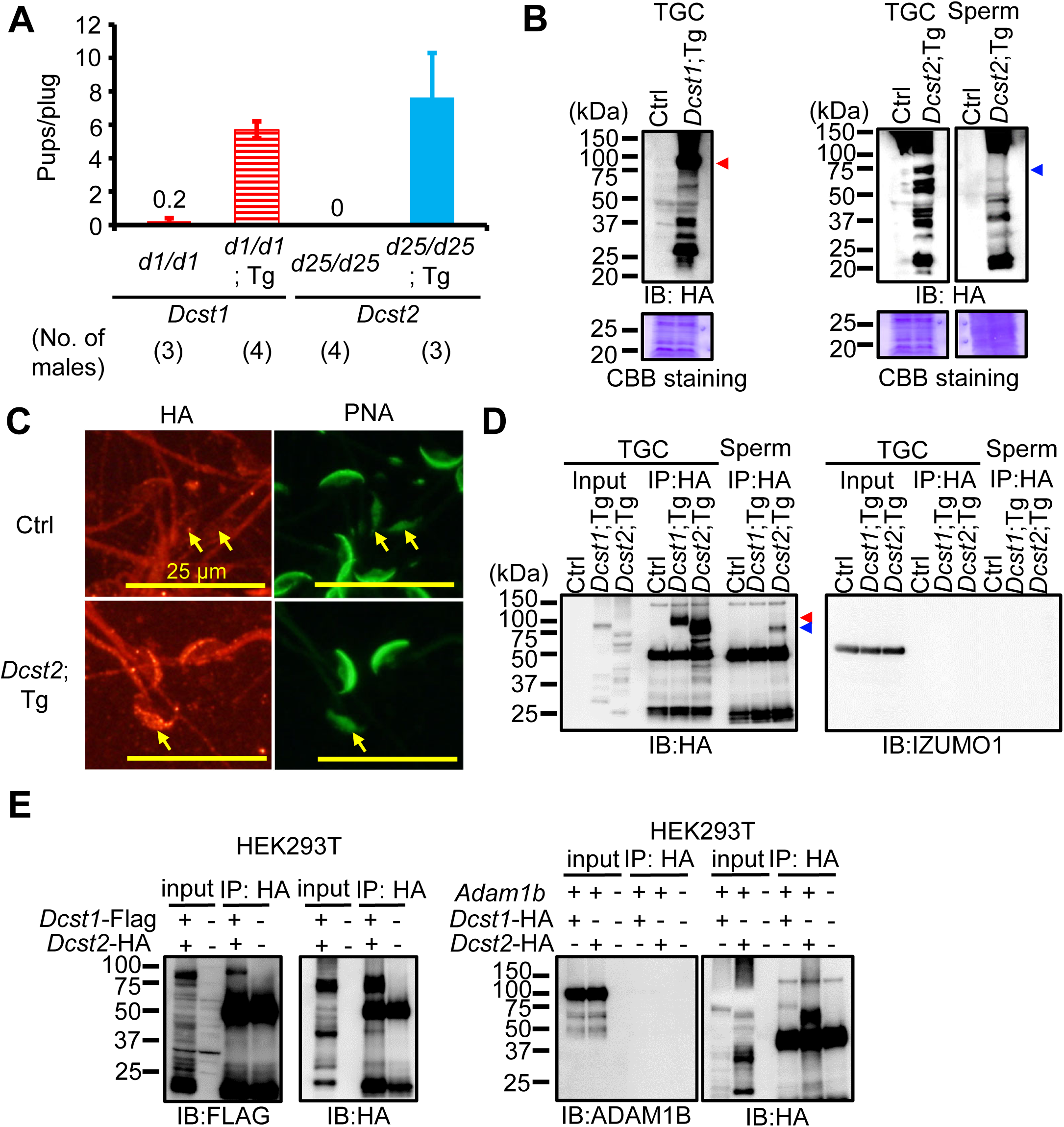
Detection of DCST1/2 in TGC and spermatozoa and interaction of DCST1/2. **A) Rescue of male fertility.** *Dcst1^d1/d1^* males with *Dcst1*-3xHA Tg insertion and *Dcst2^d25/d25^* males with *Dcst2*-3xHA Tg insertion were generated (Figure S6), and their fertility was rescued [number of plugs: 17 (*Dcst1^d1/d1^*), 25 (*Dcst1^d1/d1^*;Tg), 42 (*Dcst2^d25/d25^*), 15 (*Dcst2^d25/d25^*;Tg)]. The fecundity data in *Dcst1^d1/d1^* and *Dcst2^d25/d25^* males is replicated from Figure 1C. **B) Detection of DCST1 and DCST2 in TGC and spermatozoa.** The protein extract of TGC (100 μg) and spermatozoa (6.6×10^6 spermatozoa) was used for SDS-PAGE. The HA-tagged DCST1 and HA-tagged DCST2 were detected in TGC and spermatozoa. Total proteins in the membrane were visualized by CBB staining. Triangle marks show the expected molecular size of DCST1 (about 80 kDa) and DCST2 (about 77 kDa). **C) Localization of DCST2 in spermatozoa.** The HA-tagged DCST2 was localized in the anterior acrosome before the acrosome reaction, and then translocated to the equatorial segment in acrosome-reacted spermatozoa (arrows). PNA was used as a marker for the acrosome reaction. The fluorescence in the sperm tail was non-specific. **D) Co-IP and western blotting of the interaction between IZUMO1 and DCST1/2.** The TGC and sperm lysates from Ctrl, *Dcst1*;Tg, and *Dcst2*;Tg males were incubated with anti-HA tag antibody-conjugated magnetic beads, and then the eluted protein complex was subjected to western blotting. The HA-tagged DCST1 was detected only in the IP product from TGC, and the HA-tagged DCST2 was detected in the IP-product from TGC and spermatozoa. IZUMO1 was not detected in the co-IP products. Red and blue triangle marks show the expected molecular size of DCST1 (about 80 kDa) and DCST2 (about 77 kDa), respectively. **E) Interaction between DCST1 and DCST2 in HEK293T cells.** The protein lysate collected from HEK cells overexpressing *Dcst1*-3xFLAG and *Dcst2*-3xHA was incubated with anti-HA tag antibody-conjugated magnetic beads. The FLAG-tagged DCST1 was detected in the eluted protein complex. ADAM1B, a sperm protein that localizes to the sperm surface and is not involved in sperm-egg fusion, was used for negative control.

To reveal the localization of DCST1 and DCST2 in spermatozoa, we performed immunocytochemistry with an antibody detecting the HA epitope and peanut agglutinin (PNA) as a marker for the sperm acrosome reaction. As shown in Figure 3C, PNA in the anterior acrosome was translocated to the equatorial segment after the acrosome reaction as shown previously^29^. While HA-tagged DCST1 could rarely be observed in spermatozoa, HA-tagged DCST2 was detected within the anterior acrosome of acrosome-intact spermatozoa, and then translocated to the equatorial segment after the acrosome reaction (Figure 3C), mirroring the relocalization of IZUMO1 upon the acrosome reaction^30^. Fluorescence in the sperm tail was observed in both control and *Dcst2-HA* Tg spermatozoa, indicating that this signal in the tail was non-specific (Figure 3C).

Taking advantage of the HA tag, we performed co-immunoprecipitation (co-IP). While HA-tagged DCST1 was detected only in TGCs, HA-tagged DCST2 was detected in both TGCs and spermatozoa (Figure 3D). We could not detect IZUMO1 in these IP samples (Figure 3D), suggesting that DCST1 and DCST2 do not form a complex with IZUMO1. However, co-expression of Dcst1-3xFLAG and Dcst2-3xHA in HEK293T cells revealed the presence of a DCST1/DCST2 complexes (Figure 3E), which is in line with proposed complex formation between OCSTAMP and DCSTAMP during osteoclast fusion^31–33^.

### HEK293T cells expressing DCST1/2 and IZUMO1 bind to, but do not fuse with, ZP-free eggs

To assess whether DCST1 and DCST2 are sufficient for inducing sperm-egg fusion, we overexpressed *Dcst1*-3xFLAG, *Dcst2*-3xHA, and *Izumo1*-1D4 in HEK293T cells (Figure 4A). HEK293T cells overexpressing IZUMO1 could bind to, but not fuse with, ZP-free eggs (Figure 4B), which was consistent with previous reports^4, 9^. In contrast, HEK293T cells overexpressing only DCST1 and DCST2 failed to bind to ZP-free eggs (Figure 4B and C). Co-expression of IZUMO1 and DCST1/2 allowed the cells to bind to ZP-free eggs [4.24 ± 2.41 cells/eggs (IZUMO1), 2.01 ± 1.93 cells/eggs (DCST1/2 and IZUMO1)], but did not facilitate fusion with the oolemma (Figure 4B and C). Thus, though DCST1 and DCST2 appear to have a role in the sperm-egg fusion process, they are not sufficient to induce fusion, even in conjunction with IZUMO1.

**Figure 4.**
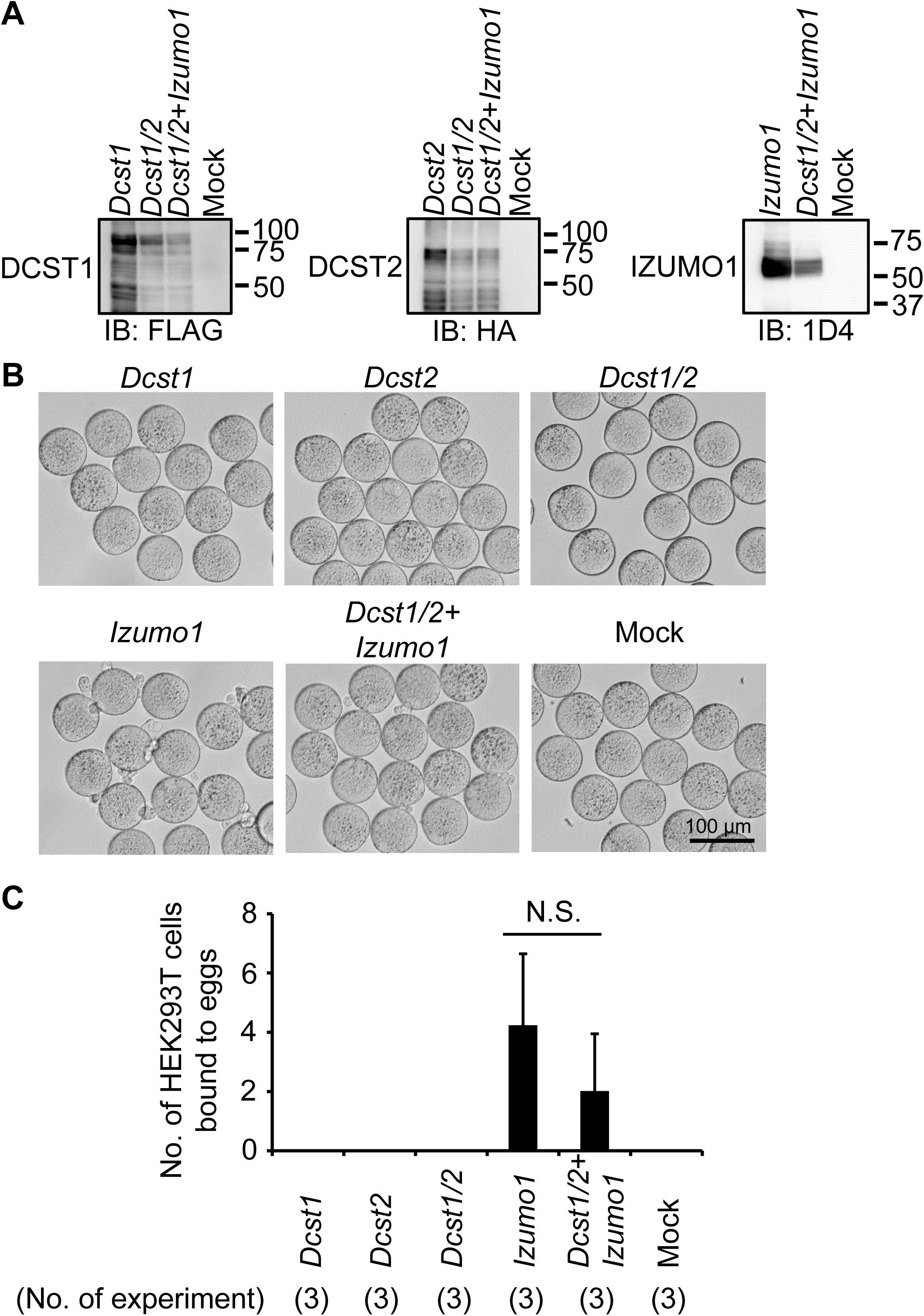
Binding assay between ZP-free eggs and HEK293T cells overexpressing *Dcst1/2*. **A) Detection of DCST1/2 and IZUMO1.** FLAG-tagged DCST1, HA-tagged DCST2, and 1D4-tagged IZUMO1 were detected in HEK293T cells overexpressing *Dcst1*-3xFLAG, *Dcst2*-3xHA, and *Izumo1*-1D4. **B and C) Observation of ZP-free eggs incubated with HEK293T cells overexpressing *Dcst1/2* and *Izumo1*.** The HEK293T cells overexpressing *Dcst1* or *Dcst2* did not attach to the oocyte membrane. Even when the HEK cells overexpressing *Dcst1/2* were used for the assay, these cells failed to bind to ZP-free eggs. The HEK293T cells overexpressing *Dcst1/2* and *Izumo1* could bind to the oocyte membrane but could not fuse with an egg. N.S.: not significant (p = 0.28).

### Sperm-expressed Dcst1/2 are also required for fertilization in zebrafish

To assess to what extent our findings in mice could be expanded among vertebrate species, we asked what the roles of DCST1/2 are in an evolutionarily distant vertebrate species, the zebrafish. The orthologous zebrafish genes *dcst1* and *dcst2* are expressed specifically in testis and arranged similarly to mouse *Dcst1/2* (Figure S7A **and** B). We therefore generated three independent KO fish lines, *dcst1*^-/-^, *dcst2^-/-^*, and *dcst1/2*^-/-^, by CRISPR/Cas9-mediated mutagenesis (Figure S7B **and** C).

Lack of zebrafish Dcst2 alone or in combination with Dcst1 caused complete sterility in males, whereas lack of Dcst1 alone led to severe subfertility [5.5 ± 3.6% fertilization rate (*dcst1*^-/-^, 16 clutches)] (Figure 5A). The fertility of heterozygous males and KO females, however, was comparable to wild-type control males. (Figure 5A). Thus, similar to mice, Dcst1/2 are essential for male fertility in zebrafish. For further phenotypic analyses we decided to focus on the *dcst2^-/-^* mutant (unless stated otherwise), since loss of Dcst2 on its own is sufficient to cause complete sterility.

**Figure 5:**
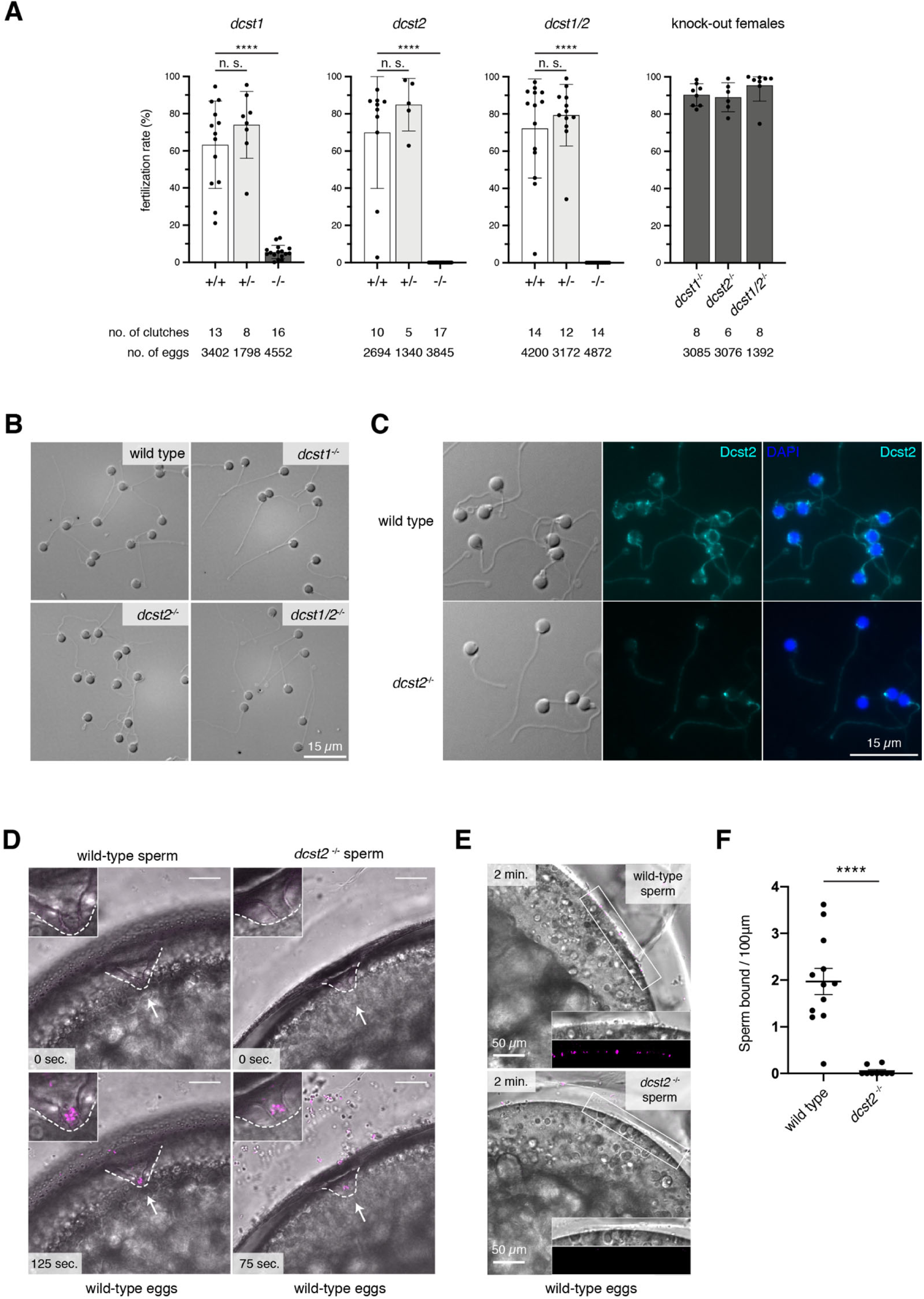
*Dcst1* and *dcst2* are essential for male fertility in zebrafish. **A) *Dcst1* and *dcst2* mutant zebrafish are male sterile.** Quantification of fertilization rates as assessed by the number of embryos that progress beyond the one-cell stage. Left three panels: Males of different genotypes [wild-type sibling (+/+; white); heterozygote sibling (+/-; light grey); homozygote sibling (-/-; dark grey)] were crossed to wild-type females; right panel: homozygous mutant females (-/-; dark grey) of the indicated genotypes were crossed to wild-type males. The number of individual clutches and the total number of eggs per genotype are indicated. Data are means ± SD; adj. ****p < 0.0001 (Kruskal-Wallis test with Dunn’s multiple-comparisons test); n. s., not significant. **B) *Dcst1* and *dcst2* mutant spermatozoa are morphologically normal.** Representative differential interference contrast images of spermatozoa from wild-type, *dcst1^-/-^*, *dcst2^-/-^*, and *dcst1/2^-/-^* fish. Scale bar: 15 µm. **C) Dcst2 localizes to the sperm head.** Immunofluorescent detection of Dcst2 protein (cyan) in permeabilized zebrafish wild-type (top) or *dcst2^-/-^* (bottom) spermatozoa using an antibody recognizing the RING-finger domain of zebrafish Dcst2. A counterstain with DAPI (blue) detects the sperm DNA in the nucleus. Scale bar: 15 µm. **D) *Dcst2* mutant spermatozoa are motile and reach the micropyle.** Images from time-lapse movies of wild-type (left) or *dcst2^-/-^* (right) spermatozoa added to wild-type eggs. Spermatozoa (magenta) were labelled with MitoTracker and added to inactivated eggs. Spermatozoa and eggs were activated by addition of water just before the start of the movie. The micropyle (white arrow), a preformed funnel in the egg coat through which the spermatozoa reach the oolemma, is outlined with a dashed white line. Top images depict the first acquired image following sperm addition: no spermatozoon has entered the micropylar area in either wild-type or mutant samples. Bottom images (125 and 75 seconds after sperm addition in wild-type and *dcst2*^-/-^ samples, respectively): spermatozoa can readily be detected within the micropylar area (inset). **E and F) *Dcst2* mutant spermatozoa are defective in stable binding to wild-type eggs.** Images from a time-lapse movie of wild-type (top) or *dcst2^-/-^* (bottom) spermatozoa added to activated and dechorionated wild-type eggs. Spermatozoa (magenta) were labelled with MitoTracker and activated at the time of addition to the eggs. Wild-type spermatozoa show clear binding to the surface of the egg (inset), while *dcst2^-/-^* spermatozoa are unable to stably bind to the oolemma (E). Binding of spermatozoa was assessed by quantifying the number of stably-bound spermatozoa in a 1-minute time window (F). ****p < 0.0001 (Mann-Whitney test).

To understand what causes the fertility defect, we first determined whether spermatozoa were produced in mutant males. *Dcst1*^-/-,^ *dcst2^-/-^*, and *dcst1/2*^-/-^ males showed normal mating behavior and produced morphologically normal spermatozoa, indicating that zebrafish Dcst1/2 are not crucial for spermatogenesis (Figure 5B). To examine where Dcst2 is localized in wild-type zebrafish spermatozoa, which lacks an acrosome, we produced antibodies against the C-terminal RING finger domain of zebrafish Dcst2. Dcst2 antibodies could detect zebrafish Dcst2 protein as determined by western blotting of wild-type and *dcst2*^-/-^ sperm lysates (Figure S7D) and immunofluorescence staining of zebrafish embryos overexpressing *dcst2(RING)-superfolder GFP (sfGFP)* mRNA (Figure S7E). Interestingly, immunofluorescence against Dcst2 strongly stained wild-type spermatozoa at the periphery of the head in punctae and occasionally the mid-piece (Figure 5C). Weaker staining of the tail region was also detected in *dcst2* KO spermatozoa, suggesting that this signal was unrelated to Dcst2. Thus, Dcst2 localizes to the periphery of the sperm head.

When added to wild-type eggs, *dcst2* KO spermatozoa were able to reach and enter the micropyle, the funnel-shaped site of sperm entry, similar to wild-type spermatozoa (Figure 5D**; Movie S3 and S4**). We therefore conclude that Dcst2 is neither required for overall sperm motility nor for spermatozoa to approach and enter the micropyle. However, in contrast to wild-type spermatozoa which remained attached to the egg (**Movie S3**), most of the entering mutant spermatozoa subsequently detached and drifted away from the micropyle (**Movie S4**), suggesting that spermatozoa lacking Dcst2 are defective in stable binding to the oolemma. We previously established an assay to assess sperm-egg binding during zebrafish fertilization^34^. Building on this assay, we used live imaging of spermatozoa and eggs to quantify the number of wild-type sperm adhered to the oolemma within a physiologically relevant time frame [1.97 ± 0.97 spermatozoa/100 µm (12 eggs)] (Figure 5E-F **and Movie S5**). Performing this assay with *dcst2* KO spermatozoa revealed that these spermatozoa were unable to adhere stably to wild-type eggs [0.05 ± 0.1 spermatozoa/100 µm (9 eggs)] (Figure 5E-F **and Movie S5**). We therefore conclude that zebrafish Dcst2 is required for stable binding of spermatozoa to the oolemma.

## Discussion

Here, we demonstrate that the testis-enriched proteins DCST1/2 are necessary for male fertility in mice and fish. These results agree with the recently reported independent findings in mice^35^ and the known requirements of *Drosophila* Sneaky and *C. elegans* SPE-42/49 in sperm for successful reproduction^21–25^, indicating that the physiological function of DCST1/2 is widely conserved among bilaterians. The conservation of DCST1/2 in sequence and function is remarkable for the otherwise rapidly evolving group of sperm-egg interacting proteins^36, 37^. Contrary to vertebrates, which have both DCST1/2 and the related DCSTAMP/OCSTAMP proteins (Figure S1A), invertebrates have solely DCST1/2.

Having established the essential role of DCST1/2 for male fertility, questions arise concerning the molecular processes in which they are involved and how they contribute to achieve fertilization. We detected mouse DCST2 at the equatorial segment of acrosome-reacted spermatozoa and zebrafish Dcst2 at the periphery of the sperm head. Given that DCST1/2 are TM proteins, these localization patterns suggest that part of the proteins are exposed on the sperm surface. Given our result that DCST1/2 form a complex, we speculate that a DCST1/2 complex in the sperm membrane either helps organize the presentation of other fusion-related sperm proteins or directly interacts with other binding-and/or fusion-relevant molecules on the oolemma. To test this hypothesis, we overexpressed DCST1 and DCST2 in IZUMO1-expressing HEK293T cells. These cells could bind to, but not fuse with, ZP-free eggs (Figure 4B-C), suggesting that DCST1/2 are not sufficient to induce fusion in a heterologous system. This observed lack of fusion could be due to the absence of other sperm-oocyte fusion-related factors (FIMP, SOF1, TMEM95, and SPACA6). Future investigation is needed to uncover the mechanism by which DCST1/2 act during fertilization.

The function of DCST1/2 in the sperm-egg fusion process differs between mice and fish: mouse DCST1/2 are required for the fusion process after sperm-egg binding (Figure 2), while zebrafish Dcst1/2 are required for sperm-egg binding (Figure 5). Given the diversity of the fertilization process across the animal kingdom, it may be that while DCST proteins are highly conserved, they may have evolved different roles to fit into the specific context of fertilization for a given species or animal group. Application of CRISPR/Cas9-mediated KO technology on a larger set of animals with divergent modes of fertilization will shed light on both the conservation and possible species-specific diversification in DCST1/2’s function in fertilization.

## Materials and Methods

### Animals

B6D2F1, C57BL/6J, and ICR mice were purchased from Japan SLC and CLEA Japan. Mice were acclimated to 12-h light/12-h dark cycle. All animal experiments were approved by the Animal Care and Use Committee of the Research Institute for Microbial Diseases, Osaka University, Japan (#Biken-AP-H30-01).

Zebrafish (*Danio rerio*) were raised according to standard protocols (28°C water temperature; 14/10-hour light/dark cycle). TLAB zebrafish served as wild-type zebrafish for all experiments, and were generated by crossing zebrafish AB stocks with natural variant TL (Tüpfel longfin) stocks. *Dcst1*^-/-^, *dcst2*^-/-^, and *dcst1/2*^-/-^ mutant zebrafish were generated as part of this study as described in detail below. All fish experiments were conducted according to Austrian and European guidelines for animal research and approved by the local Austrian authorities (animal protocol GZ: 342445/2016/12).

### Mouse sample collection

Multi-tissue expression analyses were conducted as described previously^9^. For western blotting, TGC proteins were extracted with Pierce IP lysis buffer (Thermo Fisher Scientific) (Figure 3B and D) or RIPA buffer [50 mM Tris HCl, 0.15 M NaCl, 1% Sodium deoxycholate, 0.1% SDS, 1% (vol/vol) TritonX-100, pH 7.5] containing a 1% (vol/vol) protease inhibitor mixture (Nacalai Tesque) (Figure 2C). Proteins of cauda epididymal spermatozoa were extracted with Pierce IP lysis buffer containing a 1% (vol/vol) protease inhibitor mixture (Figure 3D) or SDS sample buffer containing β-mercaptoethanol (Nacalai Tesque) (Figures 2C and 3B) as described previously^38^.

### RT-PCR for mouse multi-tissue expression analyses

Total RNA was reverse-transcribed into cDNA using a SuperScript III First-Strand Synthesis System for RT-PCR (Invitrogen). PCR was conducted with primer sets (Table S1) and KOD-Fx neo (TOYOBO). The PCR conditions were initial denaturation at 94°C for 3 minutes, denaturing at 94°C for 30 seconds, annealing at 65°C for 30 seconds, and elongation at 72°C for 30 seconds for 30 or 35 cycles in total, followed by 72°C for 2 minutes.

### Single cell RNA-seq (scRNAseq) analysis

The Median-Normalized average of *Dcst1*, *Dcst2* and fusion-related genes (*Fimp*, *Izumo1*, *Sof1*, *Spaca6*, and *Tmem95*) in spermatogenesis was examined in the published scRNAseq database^28^.

### Mouse mating test

KO male mice were caged with two B6D2F1 females for more than 1 month. After the mating period, male mice were removed from the cages, and the females were kept for another 20 days to allow them to deliver offspring. Frozen spermatozoa from *Dcst1^d1/wt^* males [B6D2-Dcst1<em2Osb> RBRC#10332, CARD#2702] and *Dcst2^d25/wt^* males [B6D2-Dcst2<em2Osb> Tg(CAG/Su9-DsRed2,Acr3-EGFP)RBGS002Osb, RBRC#11243, CARD#3047] will be available through RIKEN BRC (http://en.brc.riken.jp/index.shtml) and CARD R-BASE (http://cardb.cc.kumamoto-u.ac.jp/transgenic/).

### Mouse sperm motility and *in vitro* fertilization

Cauda epididymal spermatozoa were squeezed out and dispersed in PBS (for sperm morphology) and TYH (for sperm motility and IVF)^39^. After incubation of 10 and 120 minutes in TYH, sperm motility patterns were examined using the CEROS II sperm analysis system^40–42^. IVF was conducted as described previously^43^. Protein extracts from the remaining sperm suspension in PBS and TYH drops were used for co-IP experiments.

### Antibodies

Rat monoclonal antibodies against mouse IZUMO1 (KS64-125) and mouse SLC2A3 (KS64-10) were generated by our laboratory as described previously^44, 45^. The mouse monoclonal antibody against 1D4-tag was generated using a hybridoma cell line as a gift from Robert Molday, Ophthalmology and Visual Sciences, Centre for Macular Research, University of British Columbia, Vancouver, British Columbia, Canada^46^. Mouse monoclonal antibodies against the HA and FLAG tags were purchased from MBL (M180-3) and Sigma (F3165). The Alexa Fluor 488-conjugated Lectin PNA from *Arachis hypogaea* (peanut) was purchased from Thermo Fisher Scientific (L21409). The mouse monoclonal antibody against zebrafish Dcst2 was generated by the Max Perutz Labs Monoclonal Antibody Facility. Recombinant zebrafish Dcst2 (574-709), generated by VBCF Protein Technologies, served as the antigen. Horseradish peroxidase (HRP)-conjugated goat anti-mouse immunoglobulins (IgGs) (115-036-062) and HRP-conjugated goat anti-rat IgGs (112-035-167) were purchased from Jackson ImmunoResearch Laboratories. Fluorophore-conjugated secondary antibodies, goat anti-mouse IgG Alexa Fluor 488 (A11001), goat anti-mouse IgG Alexa Fluor 546 (A11018), goat anti-mouse IgG Alexa Fluor 594 (A11005), and goat anti-rat IgG Alexa Fluor 488 (A11006) were purchased from Thermo Fisher Scientific.

### Mouse sperm fusion assay

The fusion assay was performed as described previously^9^. To visualize IZUMO1 distribution in spermatozoa, spermatozoa after incubation of 2.5 hours in TYH drops were then incubated with the IZUMO1 monoclonal antibody (KS64-125, 1:100) for 30 minutes. Then, the spermatozoa were incubated with ZP-free eggs in TYH drops with the mixture of IZUMO1 monoclonal antibody (KS64-125, 1:100) and goat anti-rat IgG Alexa Fluor 488 (1:200) for 30 minutes. Then, the eggs were gently washed with a 1:1 mixture of TYH and FHM medium three times, and then fixed with 0.2% PFA. After washing again, IZUMO1 localization was observed under a fluorescence microscope (BZ-X700, Keyence).

### HEK293T-oocyte binding assay

Mouse *Dcst1* ORF-3xFLAG, mouse *Dcst2* ORF-3xHA, mouse *Izumo1* ORF-1D4 with a Kozak sequence (gccgcc) and a rabbit polyadenylation [poly (A)] signal were inserted under the CAG promoter. These plasmids (0.67 μg/each, total 2 μg) were transfected into HEK293T cells using the calcium phosphate-DNA coprecipitation method^47^. After 2 days of transfection, these cells were resuspended in PBS containing 10 mM (ethylenedinitrilo)tetraacetic acid. After centrifugation, the cells were washed with PBS, and then incubated with ZP-free eggs. After 30 minutes and then more than 6 hours of incubation, the attached and fused cell numbers were counted under a fluorescence microscope (BZ-X700, Keyence) and an inverted microscope with relief phase contrast (IX73, Olympus). Proteins were extracted from the remaining HEK293T cells with a lysis buffer containing Triton-X 100 [50 mM NaCl, 10 mM TrisꞏHCl, 1% (vol/vol) Triton-X 100 (Sigma Aldrich), pH 7.5] containing 1% (vol/vol) protease inhibitor mixture, and then used for western blotting and co-IP.

### Co-IP

Protein extracts [1 mg (TGC), 95∼105 μg (spermatozoa), and 200 μg (HEK293T)] were incubated with anti-HA antibody coated Dynabeads Protein G for immunoprecipitation (10009D, Thermo Fisher Scientific) for 1 hour at 4°C. After washing with a buffer (50 mM Tris-HCl, 150 mM NaCl, 0.1% Triton X-100, and 10% Glycerol, pH7.5), protein complexes were eluted with SDS sample buffer containing β-mercaptoethanol (for western blotting).

### Western blotting

Before SDS-PAGE, samples were mixed with sample buffer containing β-mercaptoethanol^38^, and boiled at 98°C for 5 minutes. For mouse samples, the polyvinylidene difluoride (PVDF) membrane was treated with Tris-buffered saline (TBS)-0.1% Tween20 (Nacalai Tesque) containing 10% skim milk (Becton Dickinson and Company) for 1 hour, followed by the primary antibody [IZUMO1, SLC2A3, HA, and FLAG (1:1,000), 1D4 (1:5,000)] for 3 hours or overnight. After washing with TBST, the membrane was treated with secondary antibodies (1:1,000). For zebrafish samples, after wet transfer onto nitrocellulose, total protein was visualized by Ponceau staining before blocking with 5% milk powder in TBST. The primary antibody [mouse anti-zebrafish-Dcst2 (1:50 in blocking buffer)] was incubated overnight at 4°C. The membrane was washed with TBST before secondary antibody incubation for 1 hour. The HRP activity was visualized with ECL prime (BioRad) and Chemi-Lumi One Ultra (Nacalai Tesque) (for mouse) or ChemiDoc (BioRad) (for zebrafish). Then, the total proteins on the membrane were visualized with Coomassie Brilliant Blue (CBB) (Nacalai Tesque).

### Immunocytochemistry

After 3-hour incubation of mouse spermatozoa in TYH drops, the spermatozoa were washed with PBS. The spermatozoa suspended with PBS were smeared on a slide glass, and then dried on a hotplate. The samples were fixed with 1% PFA, followed by permeabilization with Triton-X 100. The spermatozoa were blocked with 10% goat serum (Gibco) for 1 hour, and then incubated with a mouse monoclonal antibody against HA tag (1:100) for 3 hours or overnight. After washing with PBS containing 0.05% (vol/vol) Tween 20, the samples were subjected to the mixture of a goat anti-mouse IgG Alexa Fluor 546 (1:300) and Alexa Fluor 488-conjugated Lectin PNA (1:2,000) for 1 hour. After washing again, the samples were sealed with Immu-Mount (Thermo Fisher Scientific) and then observed under a phase contrast microscope (BX-50, Olympus) with fluorescence equipment.

Zebrafish spermatozoa were fixed with 3.7% formaldehyde immediately after collection at 4°C for 20 minutes. Spermatozoa were spun onto a SuperFrost Ultra Plus slide (Fisher Scientific) with a CytoSpin 4 (Thermo Fisher Scientific) at 1,000 rpm for 5 minutes followed by permeabilization with ice-cold methanol for 5 minutes. The slide was washed with PBS before blocking with 10% normal goat serum (Invitrogen) and 40 µg/mL BSA in PBST for 1 hour and then incubated with the mouse anti-zebrafish-Dcst2 antibody (1:25) overnight at 4°C. After washing with PBST, the slide was incubated with goat anti-mouse IgG Alexa Fluor 488 (1:100) for 2 hours before washing with PBST once more. After mounting using VECTASHIELD Antifade with DAPI (Vector Laboratories), spermatozoa were imaged with an Axio Imager.Z2 microscope (Zeiss) with an oil immersion 63x/1.4 Plan-Apochromat DIC objective.

### Fertility assessment of adult zebrafish

The evening prior to mating, the fish assessed for fertility and a TLAB wild-type fish of the opposite sex were separated in breeding cages. The next morning, the fish were allowed to mate. Eggs were collected and kept at 28°C in E3 medium (5 mM NaCl, 0.17 mM KCl, 0.33 mM CaCl_2_, 0.33 mM MgSO_4_, 10^-5^% Methylene Blue). The rate of fertilization was assessed approximately 3 hours post-laying. By this time, fertilized embryos have developed to ∼1000-cell stage embryos, while unfertilized eggs resemble one-cell stage embryos. Direct comparisons were made between siblings of different genotypes (wild-type, heterozygous mutant, homozygous mutant).

### Collection of zebrafish eggs and spermatozoa

Un-activated zebrafish eggs and spermatozoa were collected following standard procedures^48^. The evening prior to sperm collection, male and female zebrafish were separated in breeding cages (one male and one female per cage).

To collect mature, un-activated eggs, female zebrafish were anesthetized using 0.1% w/v tricaine (25x stock solution in dH_2_O, buffered to pH 7.0-7.5 with 1 M Tris pH 9.0). After being gently dried on a paper towel, the female was transferred to a dry petri dish, and eggs were carefully expelled from the female by applying mild pressure on the fish belly with a finger and stroking from anterior to posterior. The eggs were separated from the female using a small paintbrush, and the female was transferred back to the breeding cage filled with fish water for recovery.

To collect wild-type or mutant spermatozoa, male zebrafish were anesthetized using 0.1% tricaine. After being gently dried on a paper towel, the male fish was placed belly-up in a slit in a damp sponge under a stereomicroscope with a light source from above. Spermatozoa were collected into a glass capillary by mild suction while gentle pressure was applied to the fish’s belly. Spermatozoa were stored in ice-cold Hank’s saline (0.137 M NaCl, 5.4 mM KCl, 0.25 mM Na_2_HPO_4_, 1.3 mM CaCl_2_, 1 mM MgSO_4_, and 4.2 mM NaHCO_3_). The male was transferred back to the breeding cage containing fish water for recovery. For western blot analysis, spermatozoa from 3 males was sedimented at 800 x *g* for 5 minutes. The supernatant was carefully replaced with 25 µL RIPA buffer [50 mM Tris-HCl (pH 7.5), 150 mM NaCl, 1 mM MgCl_2_, 1% NP-40, 0.5% sodium deoxycholate, 1X complete protease inhibitor (Roche)] including 1% SDS and 1 U/µL benzonase (Merck). After 10 minutes of incubation at RT, the lysate was mixed and sonicated 3 times for 15 seconds of 0.5-second pulses at 80% amplitude (UP100H, Hielscher) interspersed by cooling on ice.

### Zebrafish sperm approach and binding assays

#### Imaging of zebrafish sperm approach

Spermatozoa were squeezed from 2–4 wild-type and mutant male fish and kept in 150 µL Hank’s saline containing 0.5 µM MitoTracker Deep Red FM (Molecular Probes) for >10 minutes on ice. Un-activated, mature eggs were obtained by squeezing a wild-type female. To prevent activation, eggs were kept in sorting medium (Leibovitz’s medium, 0.5 % BSA, pH 9.0) at RT. The eggs were kept in place using a petri dish with cone-shaped agarose molds (1.5% agarose in sorting medium) filled with sorting medium. Imaging was performed with a LSM800 Examiner Z1 upright system (Zeiss) with a 20x/1.0 Plan-Apochromat water dipping objective. Before sperm addition, sorting media was removed and 1 mL of E3 medium was carefully added close to the egg. 5-10 µL of stained spermatozoa was added as close to the egg as possible during image acquisition. The resulting time-lapse movies were analyzed using FIJI.

#### Imaging and analysis of zebrafish sperm-egg binding

Spermatozoa was squeezed from 2–4 wild-type and mutant male fish and kept in 100 µL Hank’s saline + 0.5 µM MitoTracker Deep Red FM on ice. Un-activated, mature eggs were squeezed from a wild-type female fish and activated by addition of E3 medium. After 10 minutes, 1-2 eggs were manually dechorionated using forceps and transferred to a cone-shaped imaging dish with E3 medium. After focusing on the egg plasma membrane, the objective was briefly lifted to add 2-10 µL of stained spermatozoa (approximately 200,000-250,000 spermatozoa). Imaging was performed with a LSM800 Examiner Z1 upright system (Zeiss) using a 10x/0.3 Achroplan water dipping objective. Images were acquired until spermatozoa were no longer motile (5 minutes). To analyze sperm-egg binding, stably-bound spermatozoa were counted. Spermatozoa were counted as bound when they remained in the same position for at least 1 minute following a 90-second activation and approach time window. Data was plotted as the number of spermatozoa bound per 100 µm of egg membrane for one minute.

### Statistical analyses

All values are shown as the mean ± SD of at least three independent experiments. Statistical analyses were performed using the two-tailed Student’s t-test, Mann-Whitney U-test, and Steel-Dwass test after examining the normal distribution and variance (Figure 2B and E, Figure 4C, Figure S4B**, and** Figure S5B). For zebrafish data, statistical analyses were performed in GraphPad Prism 7 software.

## Data availability statement

RNA-seq data reported here (zebrafish adult tissues) were deposited at the Gene Expression Omnibus (GEO) and are available under GEO acquisition number GSE171906. The authors declare that the data that support the findings of this study are available from the corresponding authors upon request.

## ACKNOWLEDGMENTS

We thank Natsuki Furuta, Eri Hosoyamada, Naoko Nagasawa, and the Biotechnology Research and Development (nonprofit organization) for excellent technical assistance; Alexander Schleiffer for phylogenetic analyses; Carina Pribitzer for RNA-Seq library preparation for zebrafish adult tissues; Mirjam Binner, Anna Kogan and the animal facility personnel from the IMP for help with genotyping and taking excellent care of zebrafish; Karin Aumayr and her team of the biooptics facility at the Vienna BioCenter (VBC) for support with microscopy; the Protein Technology Facility and the Next Generation Sequencing Facility at Vienna BioCenter Core Facilities (VBCF) for recombinant protein expression and zebrafish adult tissue RNA-Seq, respectively; the Max Perutz Labs Monoclonal Antibody Facility for generating anti-zebrafish Dcst2 antibodies; the entire Pauli lab for fruitful discussions, and Ms Ferheen Abbasi for critical reading of the manuscript. This work was supported by Ministry of Education, Culture, Sports, Science and Technology/Japan Society for the Promotion of Science KAKENHI (Grants-in-Aid for Scientific Research) Grants JP18K14612 and JP20H03172 to T.N., JP15H05573, JP16KK0180, JP20KK0155, and JP21H02397 to Y.F., JP19J21619 to S.O., and JP19H05750, JP21H04753 to M.I.; Japan Agency for Medical Research and Development Grant JP21gm5010001 to M.I.; Takeda Science Foundation grants to T.N., Y.F. and M.I.; Mochida Memorial Foundation for Medical and Pharmaceutical Research grant to Y.F.; The Sumitomo Foundation Grant for Basic Science Research Projects to Y.F.; Senri Life Science Foundation grant to Y.F.; Intramural Research Fund (grants 30-2-5 and 31-6-3) for Cardiovascular Diseases of National Cerebral and Cardiovascular Center to Y.F.; Eunice Kennedy Shriver National Institute of Child Health and Human Development Grants R01HD088412 and P01HD087157 to M.I.; and the Bill & Melinda Gates Foundation (grant INV-001902 to M.I.). Work in the Pauli lab has been supported by the FWF START program (Y 1031-B28 to A.P.), the HFSP Career Development Award (CDA00066/2015 to A.P.), a HFSP Young Investigator Award to A.P., EMBO-YIP funds to A.P., a Boehringer Ingelheim Fonds (BIF) PhD fellowship to A.B., a HFSP postdoctoral fellowship to V.E.D., and a DOC PhD student fellowship from the Austrian Academy of Sciences to K.R.G. The IMP receives institutional funding from Boehringer Ingelheim and the Austrian Research Promotion Agency (Headquarter grant FFG-852936).

## Author Contributions

T.N., A.P. and M.I. conceptualized research; T.N., A.B., Y.F., K.R.G., C.E., V.E.D., S.O., S.B., M.K., and K.P. performed research; T.N., A.B., Y.F., K.R.G., C.E., V.E.D., S.B., K.P., and M.I. analyzed data; L.E.C.Q. performed analysis of RNA-Seq data; and T.N., A.B., K.R.G., V.E.D., A.P. and M.I. wrote the paper.

## Supplementary information

### Phylogenetic tree of the DC-STAMP-like containing proteins and analysis of DCST1/2 protein conformation

Sequences were collected in an HMM search using the PFAM DC-STAMP model against NCBI-nr protein or UniProt reference proteomes databases applying highly significant E-value thresholds (<1e-10), selected for a wide taxonomic range, and aligned with mafft (-linsi, v7.427)^1–3^. A region conserved within DCST1, DCST2, DCSTAMP and OCSTAMP orthologues (covering Homo sapiens DCST1 46-599) was extracted with Jalview^4^. A maximum-likelihood phylogenetic tree was reconstructed with IQ-TREE version 2.1.3 using the “Q.plant+I+G4” model selected by ModelFinder and branch support obtained with the ultrafast bootstrap method v2^5–7^. The visualization was done with iTOL v6^8^. Branches supported by ultrafast bootstrap values (>=95%) were marked with a blue dot. Sequence accessions were added next to the species names.

The alignment of mouse and zebrafish DCST1 and DCST2 amino acid sequences was performed with Clustal Omega (https://www.ebi.ac.uk/Tools/msa/clustalo/) and visualized with JalView (https://www.jalview.org/)^4^. The DC-STAMP-like protein domain prediction was derived from pfam protein domains (http://pfam.xfam.org/). Transmembrane helices were predicted with TMHMM (http://www.cbs.dtu.dk/services/TMHMM/) and Phobius (https://phobius.sbc.su.se/)9,10.

### Generation of *Dcst1* and *Dcst2* mutant mice

*Dcst1* and *Dcst2* mutant mice were generated using 4 guide RNAs (Table S2) as described previously^11–13^. Genotyping PCR was conducted with primer sets (Table S2) and KOD-Fx neo. The PCR condition was 94 °C for 3 minutes, denaturing at 94 °C for 30 seconds, annealing at 55°C (for *Dcst2* indel mutants) or 65°C (for *Dcst1* indel and *Dcst2* deletion mutants) for 30 seconds, and elongation 72°C for 30 seconds for 35 or 40 cycles in total, followed by 72°C for 2 minutes.

### Generation of Tg mice

Sequences of *Dcst1* cDNA-3xHA tag and *Dcst2* cDNA-3xHA with a rabbit poly A signal were inserted under mouse *Clgn* promoter. The linearized DNA was injected into the pronuclear of zygotes, and the injected eggs were transferred into the ampulla of pseudopregnant females. Genotyping PCR was conducted with primer sets (Table S3) and KOD-Fx neo. The PCR condition was 94 °C for 3 minutes, denaturing at 94 °C for 30 seconds, annealing at 65°C for 30 seconds, and elongation 72°C for 1 minute (for *Dcst1*-3xHA), and 2 minutes (for *Dcst2*-3xHA) for 30 cycles in total, followed by 72°C for 2 minutes.

### Morphology and histological analysis of a mouse testis

The testicular weight and body weight of *Dcst1^d1/wt^*, *Dcst1^d1/d1^*, *Dcst2^d25/wt^*, and *Dcst2^d25/25^* males (8-24 weeks old) were measured. The testis was fixed with Bouin’s fluid (Polysciences) at 4°C overnight. Fixed samples were dehydrated by increasing the ethanol concentration, and then embedded with paraffin. Paraffin sections (5 μm) were stained with 1% periodic acid solution (Wako) for 10 minutes. Then, these samples were counterstained with Mayer hematoxylin solution for 3 to 5 min, dehydrated in increasing ethanol concentrations, and finally mounted in Entellan new (Merck).

### CRISPR/Cas9-mediated zebrafish mutant generation

Homozygous mutants for *dcst1*, *dcst2*, and *dcst1/2* were generated in zebrafish using CRISPR/Cas9-mediated mutagenesis through the use of single guide RNAs (sgRNAs) generated as previously described^14^. SgRNAs targeting exons 2 and 3 for *dcst1* or exon 4 for *dcst2* (Table S4) were co-injected with Cas9 protein into one-cell TLAB embryos to generate the single knock-out mutants (Figure S7B). *Dcst1/2*^-/-^ zebrafish were generated in the same manner, but by injecting sgRNAs targeting exons 2 and 3 for *dcst1* in conjunction with those targeting exon 4 for *dcst2* into a *dcst1* mutant background (149 bp deletion, 14 bp insertion) to increase the chances of mutagenesis for both genes on the same allele. For all mutants, injected embryos were grown to adulthood and out-crossed to *wt* TLAB zebrafish; the offspring were then screened by PCR (Table S4) for heterozygous mutations in *dcst1*, *dcst2*, or both loci to identify founders. Siblings of fish found to carry mutations in the *dcst1* or *dcst2* locus were grown to adulthood and in-crossed to generate homozygous mutant fish. Amplicon sequencing of adult fin-clips revealed the different mutations as a 2-bp substitution combined with a 50-bp insertion and 1-bp substitution in exon 2, and 4-bp deletion in exon 3 of *dcst1*^-/-^; a 7-bp deletion in exon-4 for *dcst2*^-/-^; and a 155-bp deletion in *dcst1* exon 3 combined with a 4-bp insertion followed by a 64-bp insertion in *dcst2* exon 4 in case of the double mutant *dcst1/2*^-/-^.

Genotyping of *dcst1*^-/-^, *dcst2*^-/-^, and *dcst1/2*^-/-^ mutant fish was performed using PCR (Table S4). Detection of mutations was performed using standard agarose gel electrophoresis (wt amplicon for *dcst1*: 311 bp, wt amplicon for *dcst2*: 394 bp, *dcst1*^-/-^ amplicon: 358 bp; *dcst2*^-/-^ amplicon: 387 bp; amplicon sizes in the *dcst1/2*^-/-^ double mutant: *dcst1*^-/-^ amplicon: 156 bp; *dcst2*^-/-^ amplicon 467 bp). To confirm the presence of a truncated mRNA product for both mutants, cDNA was generated using the iSCRIPT cDNA Synthesis Kit (BioRad) from RNA isolated from mutant testis tissue. Sanger sequencing using primers within the open reading frame for both *dcst1* and *dcst2* was performed using the primers listed below (Table S4). Confirming that the mutations lead to a truncated mRNA product, the *dcst1*^-/-^ cDNA encoded only 15 amino acids with a premature termination codon, compared to 676 amino acids for wt, the *dcst2*^-/-^ cDNA encoded 183 amino acids (709 amino acids in wt), and the *dcst1/2*^-/-^ cDNA encoded 25 amino acids for *dcst1* and 170 amino acids for *dcst2* (Figure S7C).

### Generation of Dcst2 RING finger domain constructs

To clone the coding sequence of the Dcst2 RING finger domain, RNA was isolated from adult testis using the standard TRIzol (Invitrogen) protocol followed by phenol/chloroform extraction. cDNA was synthesized using the iSCRIPT cDNA Synthesis Kit (BioRad) and served as template for the amplification of the sequence underlying Dcst2 (566-709) (Dcst2-RING_F and Dcst2-RING_R). The PCR product was introduced into BamHI/EcoRI-cut *pMTB-actb2:MCS-sfGFP* (a derivative of *pMTB-actb2:H2B-Cerulean*, a kind gift from Sean Megason) by Gibson assembly^15^ to generate an in-frame fusion to sfGFP. The resulting ORF is flanked by a SP6 promoter and SV40 3’ UTR.

### mRNA injection of zebrafish embryos

TLAB embryos were collected immediately after being laid and dechorionated with pronase (1 mg/ml). Dechorionated one-cell stage embryos were injected with 100 pg mRNA and cultured at 28°C in E3 medium (5 mM NaCl, 0.17 mM KCl, 0.33 mM CaCl_2_, 0.33 mM MgSO_4_, 10^-5^% Methylene Blue). mRNA encoding Dcst2(566-709)-sfGFP was prepared by *in vitro* transcription using the mMESSAGE mMACHINE SP6 kit (Invitrogen) from a plasmid containing the SP6 promoter and SV40 3’ UTR. Shield-stage (6 hpf) embryos were fixed with 3.7% formaldehyde at 4°C overnight. Immunofluorescence staining of embryos was performed as was done for zebrafish sperm immunocytochemistry, but with goat anti-mouse IgG Alexa Fluor 594 as secondary antibody. The embryos were imaged in PBS in a watch glass with a stereomicroscope (Lumar, Zeiss) with the 0.8x Neo Lumar S objective and 5x zoom (40x magnification).

### Zebrafish adult tissue RNA-Seq

Adult zebrafish tissues (muscle, spleen, liver, intestine, heart, brain, fin, skin, testis) were dissected from adult *wt* (TLAB) zebrafish (three biological replicates for each sample). Total RNA was isolated using the standard TRIzol protocol and assessed for quality and quantity based on analysis on the Fragment Analyzer. PolyA+ RNA was selected with the poly(A) RNA Selection Kit (LEXOGEN), following the manufacturer’s instructions. Stranded cDNA libraries were generated using NEBNext Ultra Directional RNA Library Prep Kit for Illumina (New England Biolabs) and indexed with NEBNext Multiplex Oligos for Illumina (Dual Index Primer Set I) (New England Biolabs). Library quality was checked on the Fragment Analyzer and sequenced on a Illumina Hiseq 2500 with the SR100 mode. RNA-seq reads were processed according to standard bioinformatic procedures. Reads were mapped to Ensembl 102 gene models (downloaded 2021.01.25) and the GRCz11 Danio rerio genome assembly, with Hisat2 v2.1.0^16^ using the Ensembl transcriptome release 102. A custom file was generated by adding bouncer based on its position coordinates [exon = chr18:50975023-50975623 (+ strand); CDS = chr18:50975045-50975422 (+ strand)]. Quantification at the gene level (transcript per million (TPM)) was performed using Kallisto (v0.46.0)^17^. The RNA-seq data set was deposited to Gene Expression Omnibus (GEO) and is available under GEO acquisition number GSE171906. RNA-seq data of ovary and oocyte-stage samples have been published previously and are available under GEO acquisition numbers GSE111882 (testis, ovary, mature oocytes)^18^ and GSE147112 (oogenesis, mature oocytes)^19^.

---

**Figure S1.**
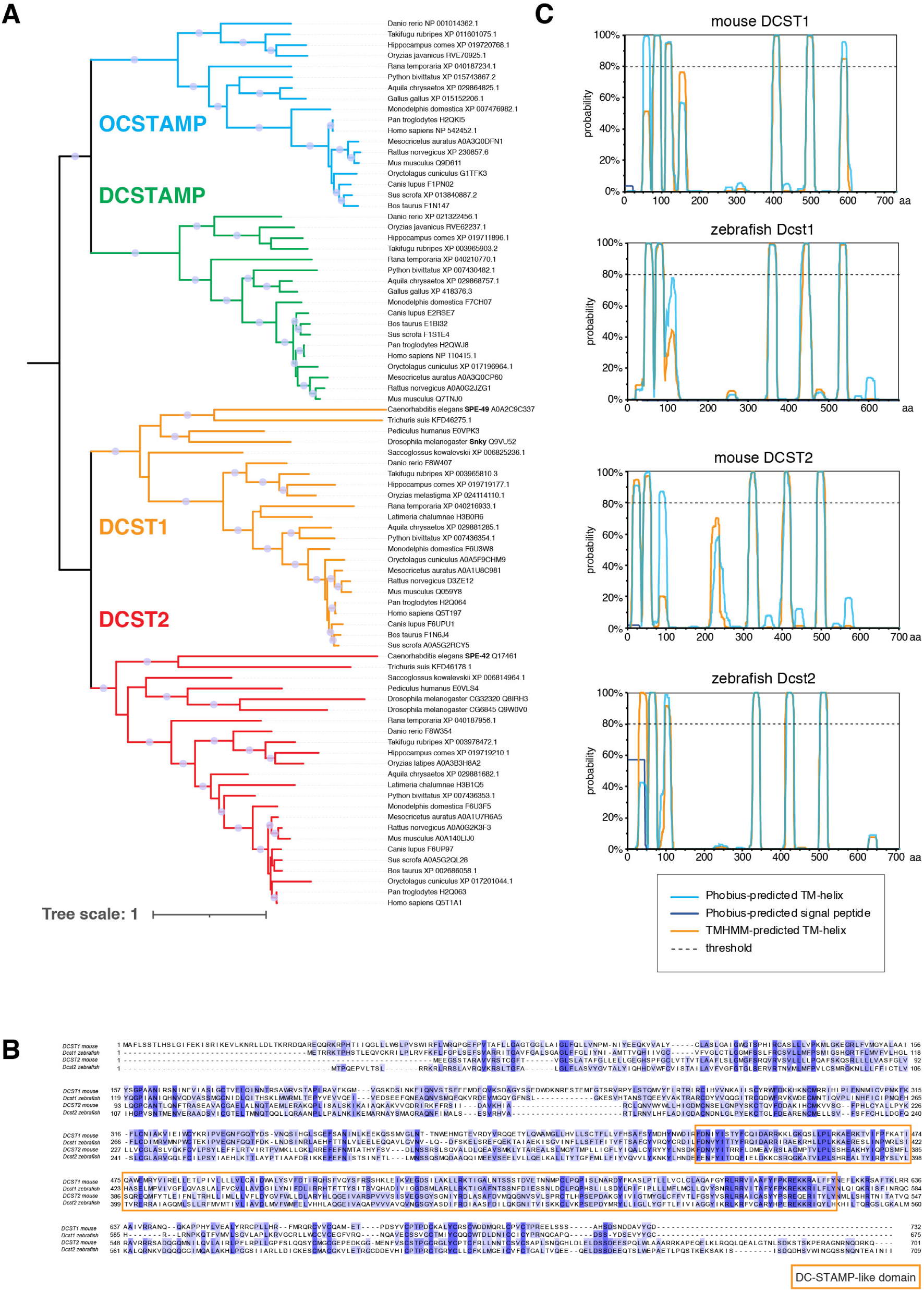
Conservation of DCST1 and DCST2 across bilaterians and predicted protein structure. **A) Phylogenetic tree of the DC-STAMP-like domain containing proteins**. A maximum-likelihood phylogenetic tree revealed an early split between DCSTAMP (green) and OCSTAMP (blue) proteins on the one hand and DCST1 (orange) and DCST2 (red) on the other hand. Common protein names for *C. elegans* SPE-42/SPE-49 and *D. melanogaster* Sneaky are included. **B) Amino acid sequence alignment between mouse and zebrafish DCST1 and DCST2**. Letters with blue shading indicate the percentage of sequence identity (dark blue: 100% identity). The DC-STAMP-like protein domain is highlighted in an orange box. **C) Predicted transmembrane helices for mouse and zebrafish DCST1 and DCST2**. Plots show predicted probabilities for transmembrane helices [TMHMM (orange) and Phobius (blue)] in mouse and zebrafish DCST1 and DCST2.

**Figure S2.**
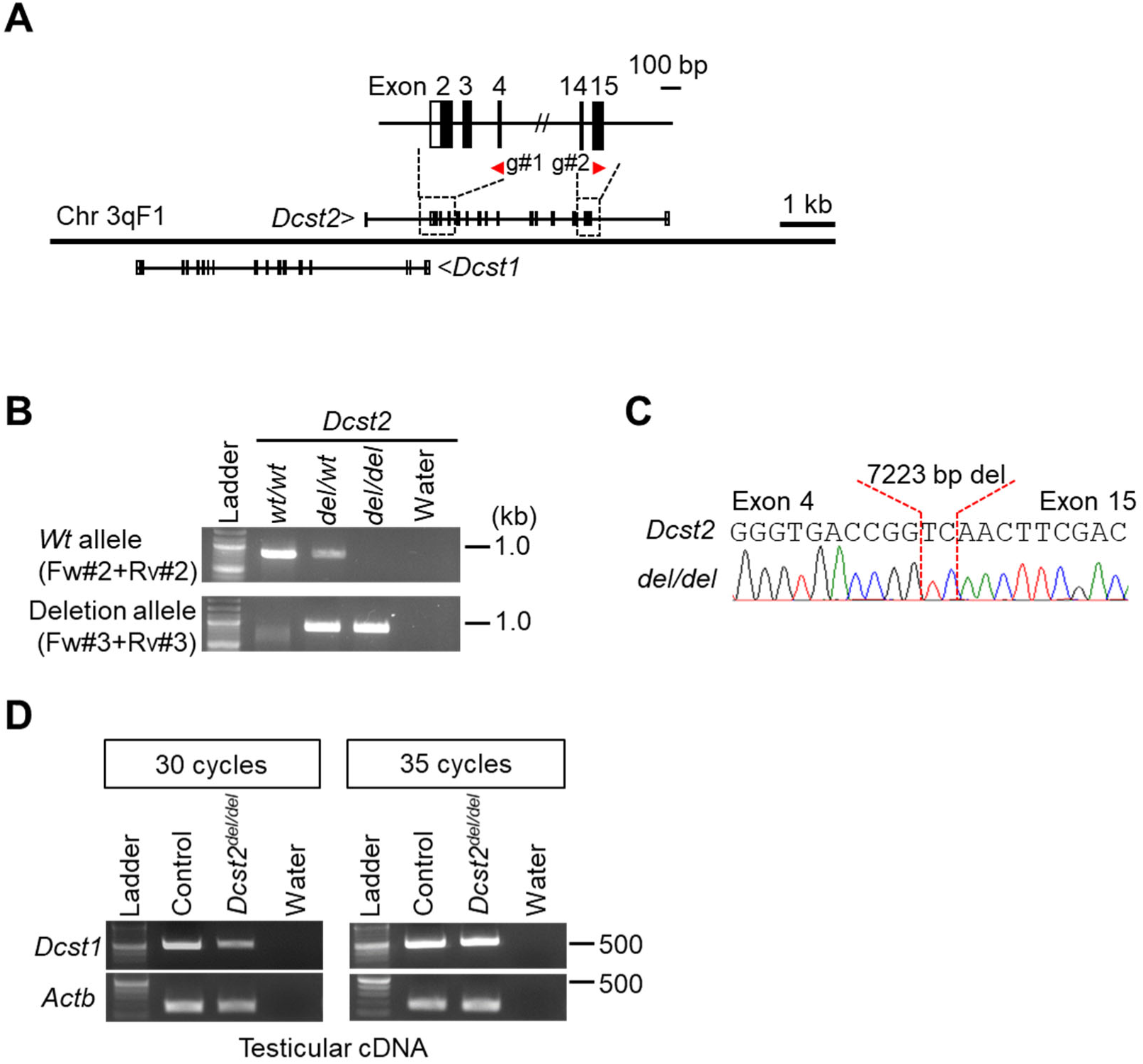
Generation of *Dcst2^del/del^* mice. **A) gRNA design.** Mouse *Dcst1* and *Dcst2* are tandemly arranged. To delete the coding region of *Dcst2*, we designed 2 gRNAs in exon4 and 15 of *Dcst2*. Black colored boxes show the coding region. **B and C) Genotyping with PCR and direct sequencing.** Four primers were used for the genotyping PCR. The amplicons were subjected to direct sequencing, and the mutant allele has a 7223 bp deletion and 2 bp insertion. **D) Detection of *Dcst1* mRNA in *Dcst2^del/del^* testis.** The expression level of *Dcst1* mRNA decreased in *Dcst2^del/del^* testis, but it still remained.

**Figure S3.**
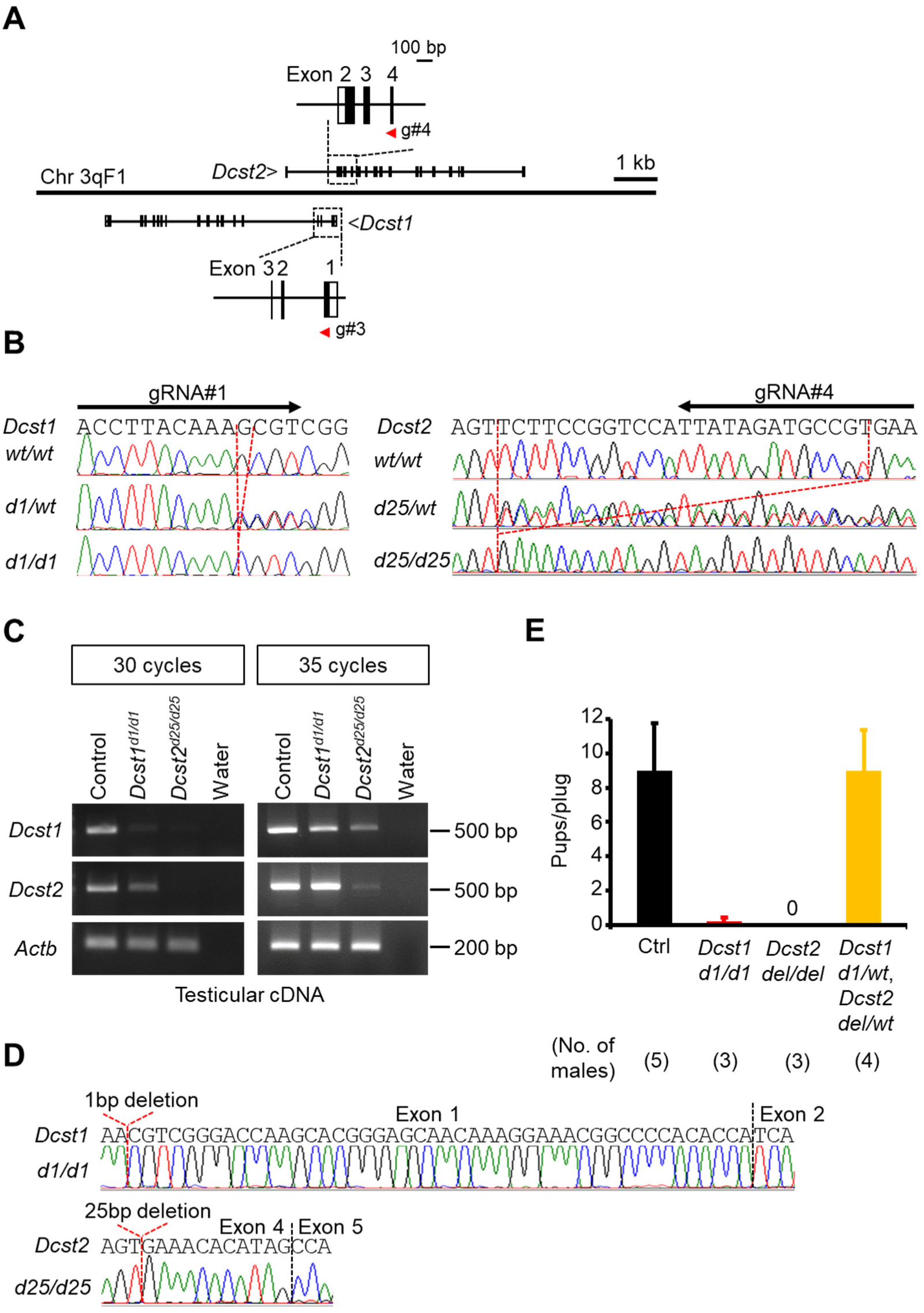
Generation of *Dcst1^d1/d1^* and *Dcst2^d25/d25^* mice. **A) gRNA design.** To generate indel mutant mice of *Dcst1* and *Dcst2*, we designed 2 gRNAs in exon1 of *Dcst1* and exon 4 of *Dcst2*. Black colored boxes show the coding region. **B) Genome sequence of *Dcst1* and *Dcst2* in *Dcst1^d1/d1^* and *Dcst2^d25/d25^* mice.** The mutant alleles of *Dcst1* and *Dcst2* have a 1-bp deletion in *Dcst1* and a 25-bp deletion in *Dcst2*, respectively. **C and D) Detection and cDNA sequencing of *Dcst1* and *Dcst2* mRNAs in *Dcst1^d1/d1^* and *Dcst2^d25/d25^* testes.** The expression level of *Dcst1* mRNA in *Dcst2^d25/d25^* testis and *Dcst2* mRNA in *Dcst1^d1/d1^* testis decreased, but still remained. *Dcst1* mRNA in *Dcst1^d1/d1^* testis and *Dcst2* mRNA in *Dcst2^d25/d25^* testis were detected (panel C), but *Dcst1* and *Dcst2* cDNA sequencing have a 1-bp and 25-bp deletion in *Dcst1* and *Dcst2*, respectively. **E) Fertility of double heterozygous (*Dcst1^d1/wt^* and *Dcst2^del/wt^*) (dHZ) males.** The fertility of dHZ males was comparable to the control [number of plugs: 24 (*Dcst1^d1/wt^*, *Dcst2^del/wt^*)]. The fecundity data in Ctrl, *Dcst1^d1/d1^* and *Dcst2^del/del^* males is from Figure 1C.

**Figure S4.**
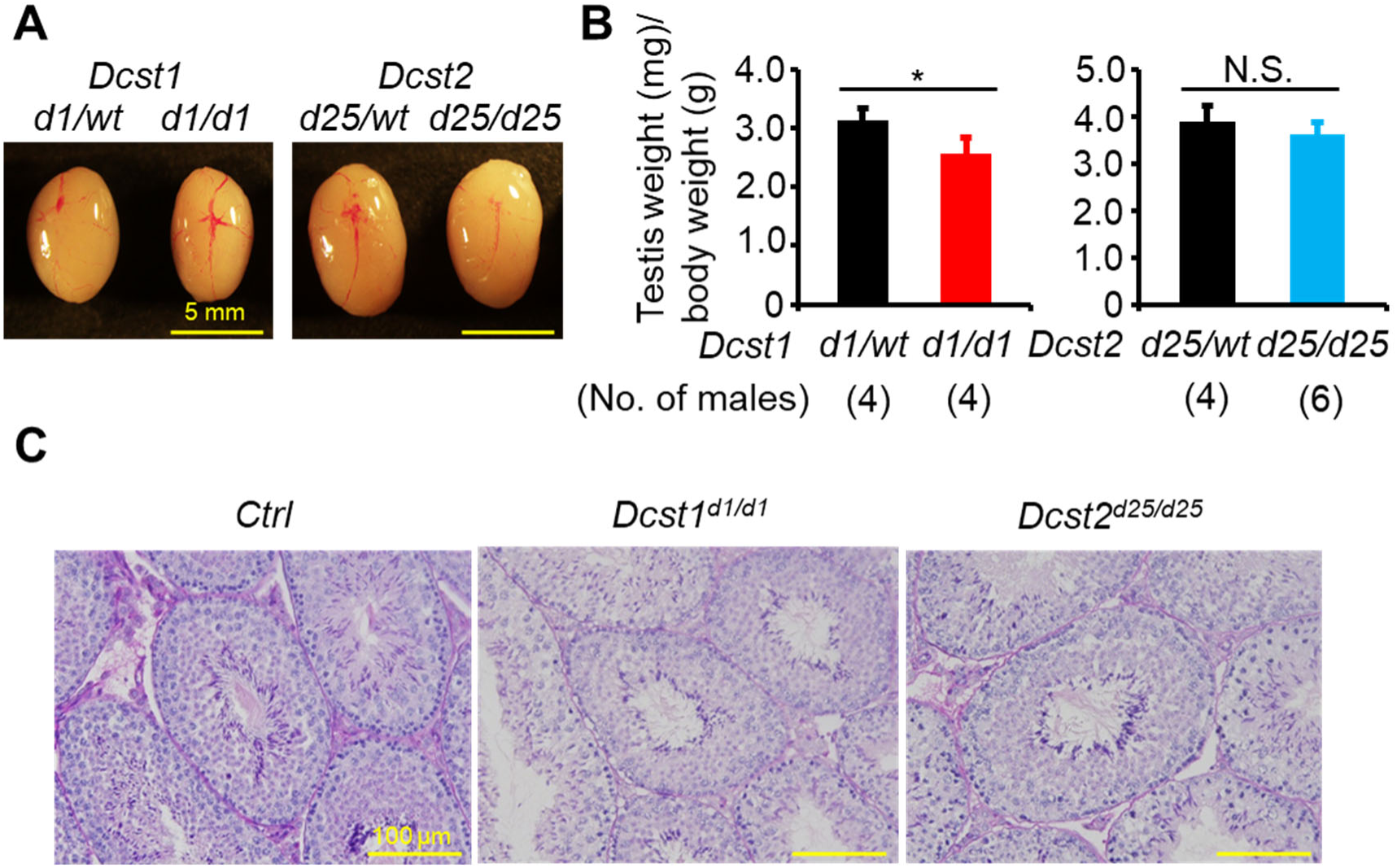
Analysis of *Dcst1^d1/d1^* and *Dcst2^d25/d25^* testes. **A) Testis morphology.** **B) Testis weight (mg) / body weight (g).** The value of *Dcst1^d1/d1^* testis slightly decreased compared with the *Dcst1^d1/wt^* testis. *: p < 0.05., N.S.: not significant (p = 0.24) **C) Histological analysis.** There was no overt abnormality in *Dcst1^d1/d1^* and *Dcst2^d25/d25^* spermatogenesis.

**Figure S5.**
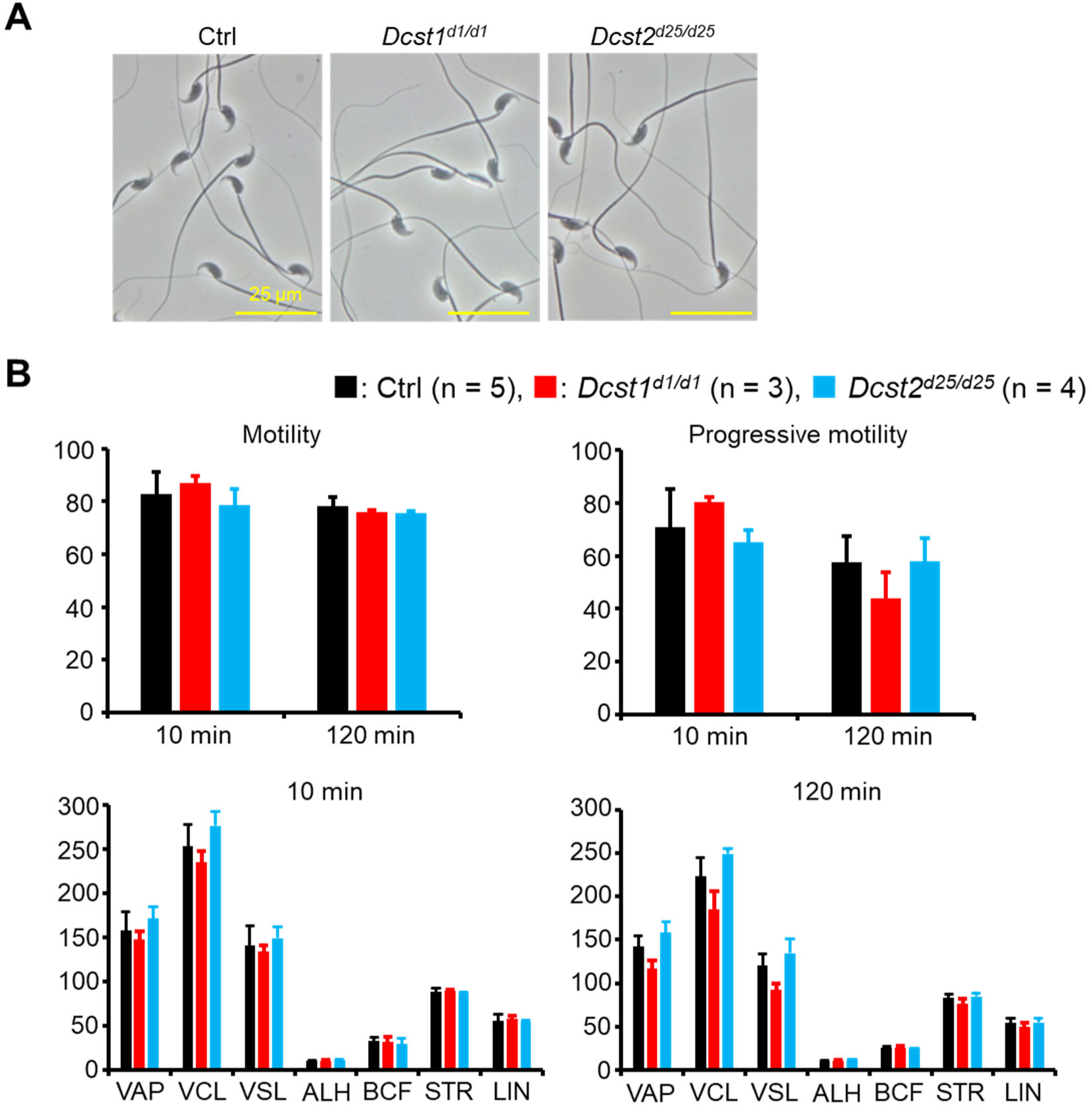
Analysis of *Dcst1* KO and *Dcst2* KO spermatozoa. **A) Sperm morphology.** **B) Sperm motility.** There was no difference in sperm motility parameters between Ctrl, *Dcst1* KO and *Dcst2* KO spermatozoa. Spermatozoa from *Dcst1^d1/wt^* and *Dcst2^d25/wt^* males were used as the control. VAP: average path velocity, VSL: straight line velocity, VCL: curvilinear velocity, ALH: amplitude of lateral head, BCF: beat cross frequency, STR: straightness of trajectory, LIN: linearity.

**Figure S6.**
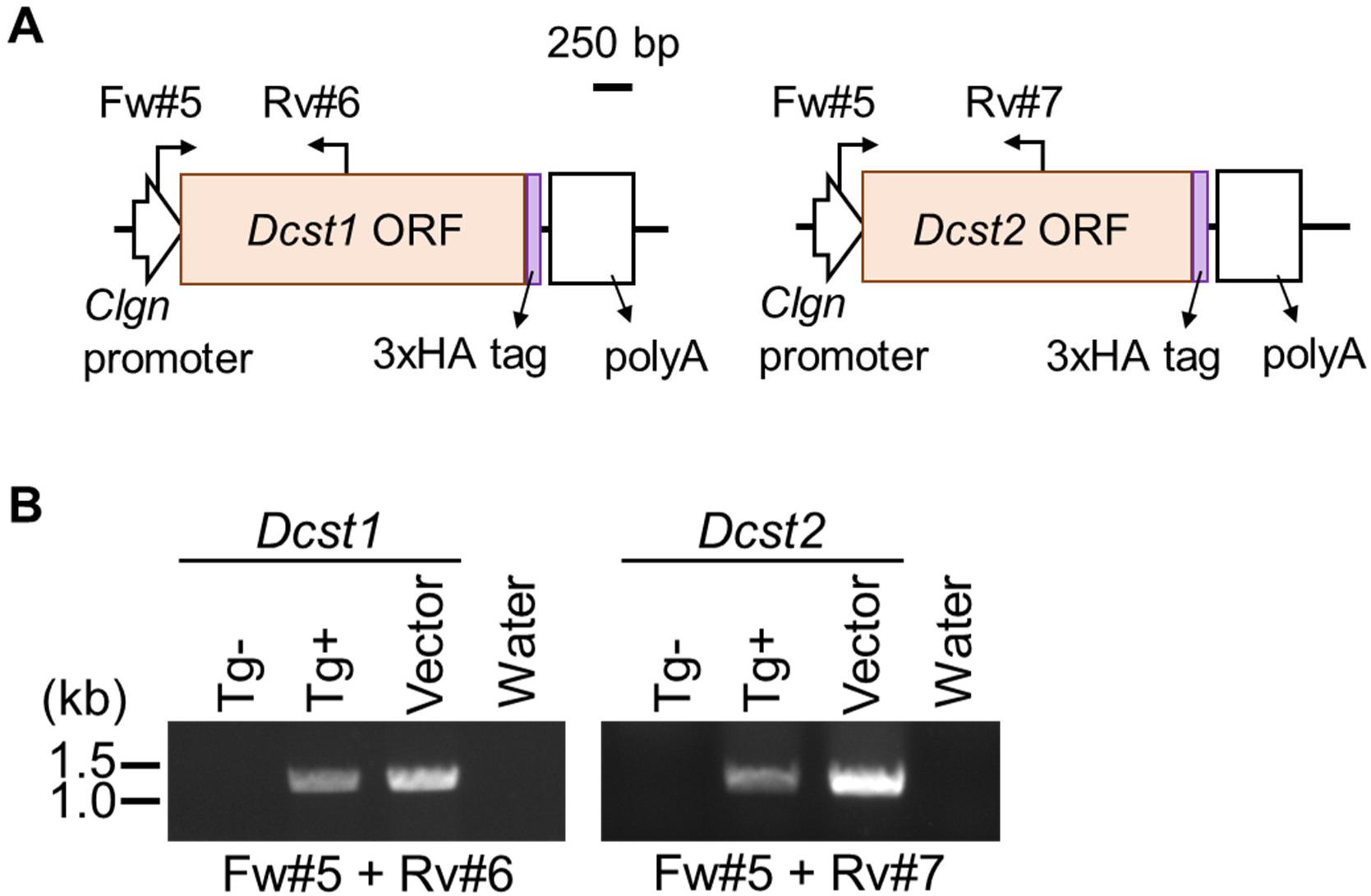
Generation of *Dcst1*-3xHA and *Dcst2*-3xHA Tg mice. **A) Schematic presentation of generating *Dcst1*-3xHA and *Dcst2*-3xHA Tg mice.** The transgenes with ORF regions of mouse *Dcst1* and *Dcst2* and 3xHA tag were expressed as a fused protein under the testis-specific Calmegin (*Clgn*) promoter. **B) Genotyping of *Dcst1*-3xHA and *Dcst2*-3xHA Tg mice.**

**Figure S7:**
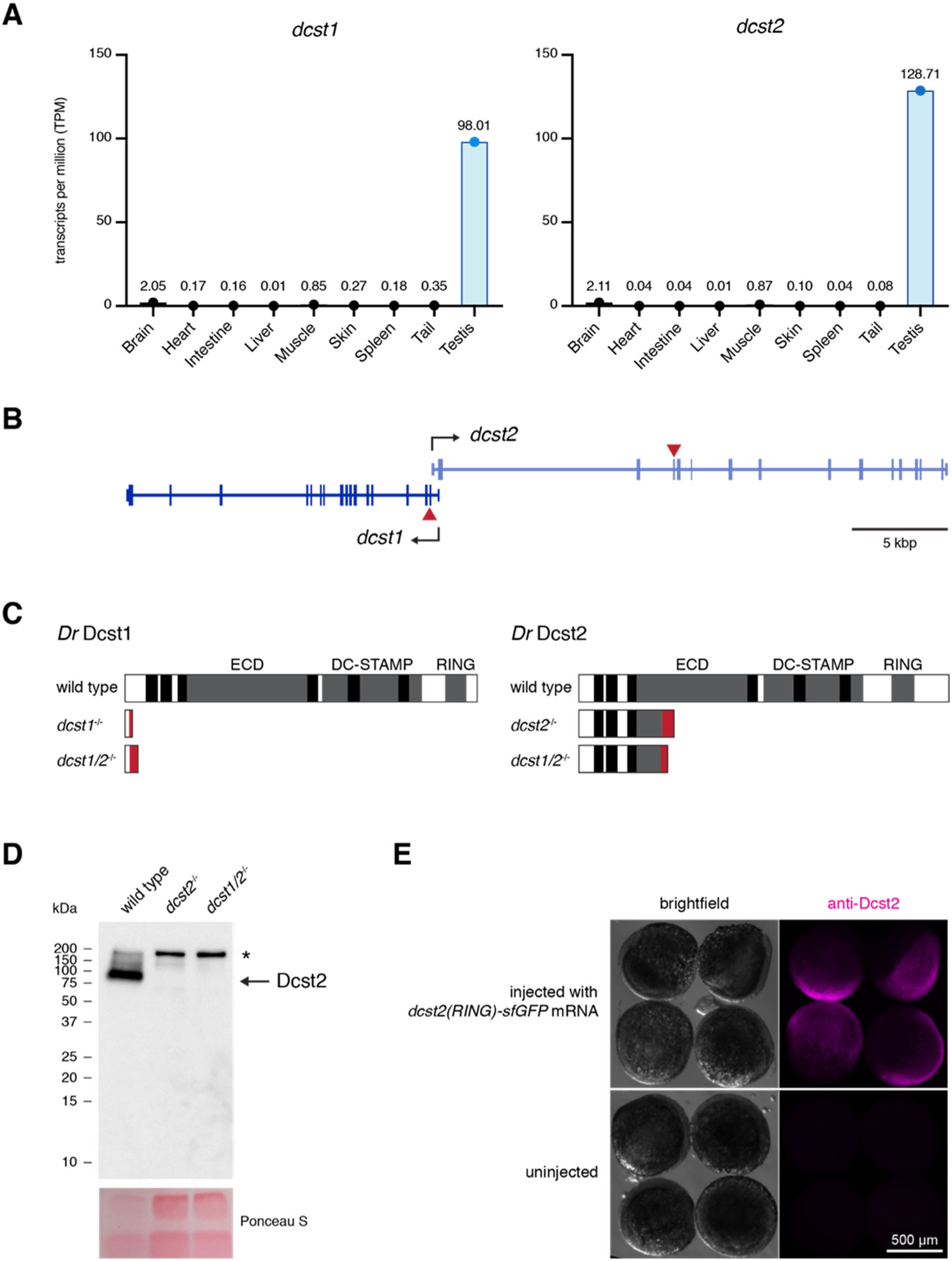
Expression of *dcst1* and *dcst2* in zebrafish. **A) *Dcst1* and *dcst2* genes are specifically expressed in adult testis in zebrafish.** RNA-Seq analysis of *dcst1* and *dcst2* gene expression levels in various adult tissues. Amongst all the tissues tested, *dcst1* and *dcst2* transcripts are strongly enriched in adult testis. The y-axis shows TPM values (transcripts per million). **B) *Dcst1* and *dcst2* gene locus in zebrafish.** *Dcst1* and *dcst2* genes overlap in their 5’ ends. The red triangles indicate the sites of the introduced mutations in *dcst1* and *dcst2*. **C) Dcst1 and Dcst2 domain organization and mutant translation.** Zebrafish Dcst1 and Dcst2 are multi-pass transmembrane proteins. Predicted transmembrane domains (black, Phobius prediction^10^), the extracellular domain (ECD), the DC-STAMP-like domain (DC-STAMP) and C_4_C_4_ RING finger domain^20^ are indicated. The *dcst1* and *dcst2* mutant alleles encode for truncated proteins. The aberrant translation caused by frameshift indels up to the premature termination codon is indicated in red. **D) Dcst2 protein is detected in wild-type spermatozoa and absent in *dcst2* mutants.** Sperm protein-extract of *wt*, *dcst2^-/-^* and *dcst1/2^-/-^* was prepared and analyzed by SDS-PAGE and western blot analysis, using an antibody recognizing the RING-domain of zebrafish Dcst2. The band corresponding to wild-type Dcst2 (predicted protein size: 83 kDa) is indicated with an arrow; the asterisk indicates an unspecific band. The Ponceau S-stained membrane is shown as a loading control. **E) The anti-Dcst2 antibody is able to detect overexpressed Dcst2 protein in zebrafish embryos by immunofluorescence.** Immunofluorescent detection of Dcst2 protein (magenta) in zebrafish embryos. Embryos were either injected at the 1-cell stage with 100 pg of *dcst2 (RING)*^20^*-sfGFP*) mRNA (top; overexpression of Dcst2 (RING)-sfGFP) or not injected (bottom; negative control). Embryos were fixed after 6 hours, followed by immuno-staining using an antibody recognizing the RING-domain of zebrafish Dcst2. Scale bar: 500 µm.

## Supplementary Movie Legends

**Movie S1. Egg observation after IVF using *Dcst1* KO spermatozoa.**

**Movie S2. Egg observation after IVF using *Dcst2* KO spermatozoa.**

**Movie S3. Wild-type sperm approach to micropyle**. Wild-type spermatozoa stained with Mitotracker Deep-Red was added to wild-type eggs and images were acquired following sperm addition.

**Movie S4. Dcst2 mutant sperm approach to micropyle**. Dcst2 mutant spermatozoa stained with Mitotracker Deep-Red was added to wild-type eggs and images were acquired following sperm addition.

**Movie S5. Dcst2 mutant spermatozoa are unable to stably bind to wild-type eggs**. Time lapse of sperm binding assay with wild-type (left) and dcst2-/- (right) spermatozoa stained with Mitotracker Deep Red and wild-type eggs. After 2 minutes following sperm addition, wild-type spermatozoa are stably bound to the oolemma while dcst2-/- mutant spermatozoa are unable to bind.

---

**Supplemental Table S1.**
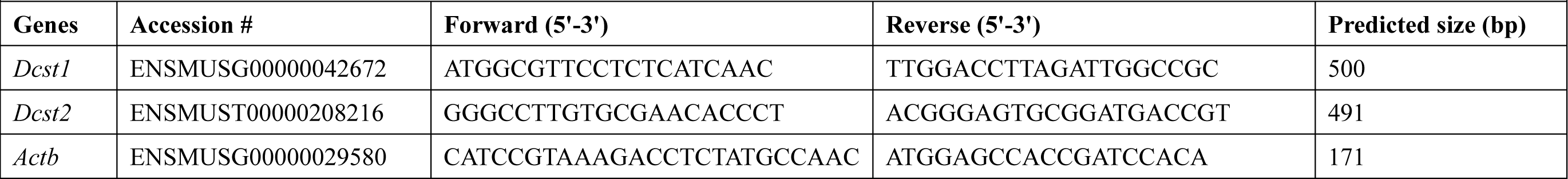
Primer sequences for the tissue expression analysis.

**Supplemental Table S2.**
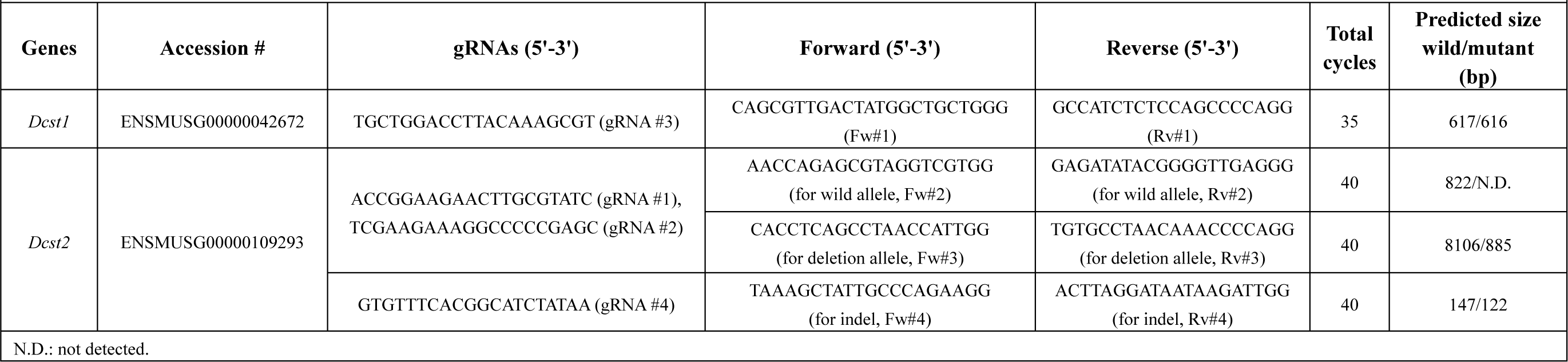
Primer sequences for the genotyping and gRNA sequences.

**Supplemental Table S3.**
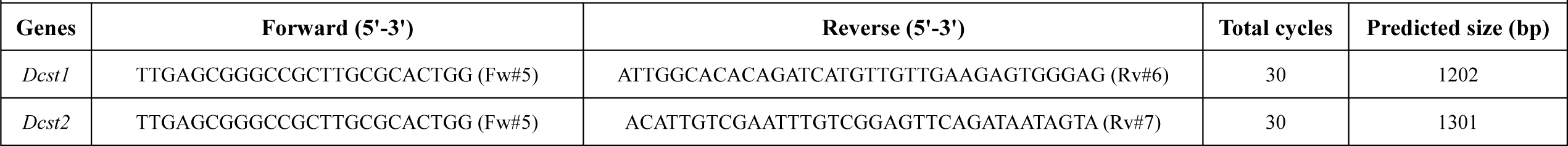
Primer sequences for the genotyping of Tg mice.

**Supplemental Table S4.**
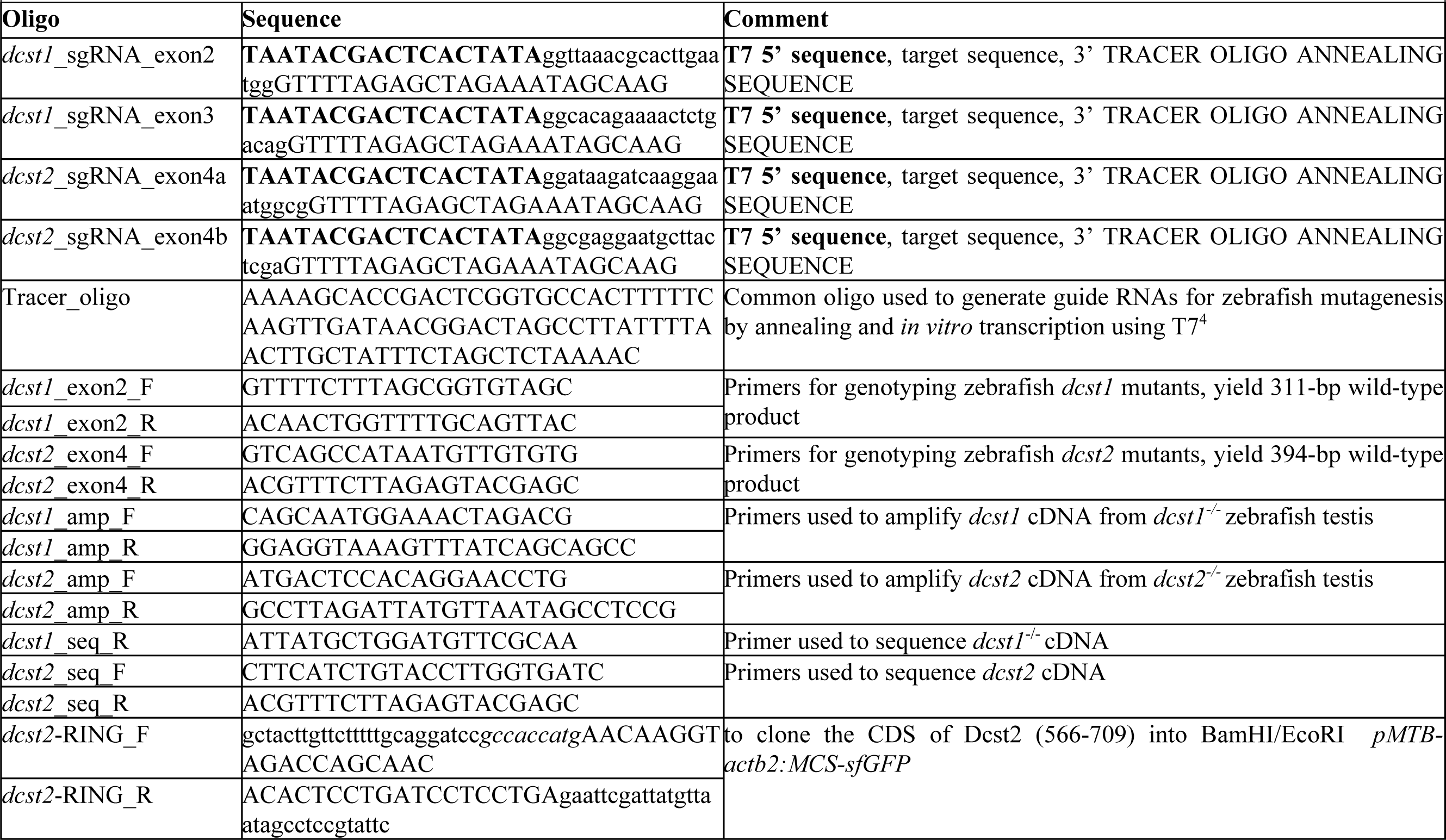
Primers for zebrafish.

